# A protein complex in the extreme distal tip of vertebrate motile cilia controls their organization, length, and function

**DOI:** 10.1101/2025.02.19.639145

**Authors:** Juyeon Hong, Chanjae Lee, Gopika Madhu, Ophelia Papoulas, Ece Atayeter, Garbiel Hoogerbrugge, Jiehong Pan, Maki Takagishi, Nadia Manzi, Daniel J. Dickinson, Amjad Horani, Steven L. Brody, Edward Marcotte, Vivek N. Prakash, Tae Joo Park, John B. Wallingford

**Affiliations:** Department of Molecular Biosciences, University of Texas at Austin, Texas, USA; Department of Biological Sciences, Ulsan National Institute of Science and Technology, Ulsan, South Korea; Department of Physics, College of Arts and Sciences, University of Miami, Coral Gables, FL 33146, USA; Division of Pulmonary and Critical Care Medicine, Department of Medicine, Washington University School of Medicine in St. Louis, USA; Dept. of Medicinal and Life Sciences, Nagoya City University, Nagoya, Japan; Department of Pediatrics, Washington University School of Medicine, Saint Louis, MO 63110, USA; Dept. of Biology, College of Arts and Sciences and Department of Marine Biology and Ecology, Rosenstiel School of Marine, Atmospheric and Earth Science, University of Miami, Coral Gables, FL, USA; Center for Genomic Integrity, Institute for Basic Science, Ulsan 44919, Republic of Korea

## Abstract

The beating of cilia on multi-ciliated cells (MCCs) is essential for normal development and homeostasis in animals. But while the structure and function of basal bodies and axonemes have received significant attention recently, the distal tips of MCC cilia remain relatively poorly defined. Here, we characterize the molecular organization of the distal tip of vertebrate MCC cilia, characterizing two distinct domains occupied by distinct protein constituents. Using frog, mouse, and human MCCs, we find that two largely uncharacterized proteins, Ccdc78 and Ccdc33 occupy a previously undefined region at the extreme distal tip, and these are required for the normal organization of all other known tip proteins. Ccdc78 and Ccdc33 each display robust microtubule-bundling activity both *in vivo* and *in vitro*, yet each is independently required for normal length regulation of MCC cilia. Moreover, loss of each protein elicits a distinct pattern of defective cilia beating and resultant fluid flow. Thus, two previously undefined proteins form a key module essential for organizing and stabilizing the distal tip of motile cilia in vertebrate MCCs. We propose that these ill-defined proteins represent potential disease loci for motile ciliopathies.

## Introduction

Motile cilia are microtubule-based organelles conserved across the eukaryotes and play important roles in development and homeostasis. Motile cilia on multi-ciliated cells (MCCs) generate fluid flow to clear mucus in the airways, to transport gametes in the reproductive tracts, and to move cerebrospinal fluid in the central nervous system (Fig. 1A)^1,2^. Dysfunction of ciliary motility in MCCs causes primary ciliary dyskinesia (PCD), an incurable genetic disease characterized by chronic respiratory tract infection leading to bronchiectasis as well as infertility and hydrocephalus^3–5^.

**Figure 1.**
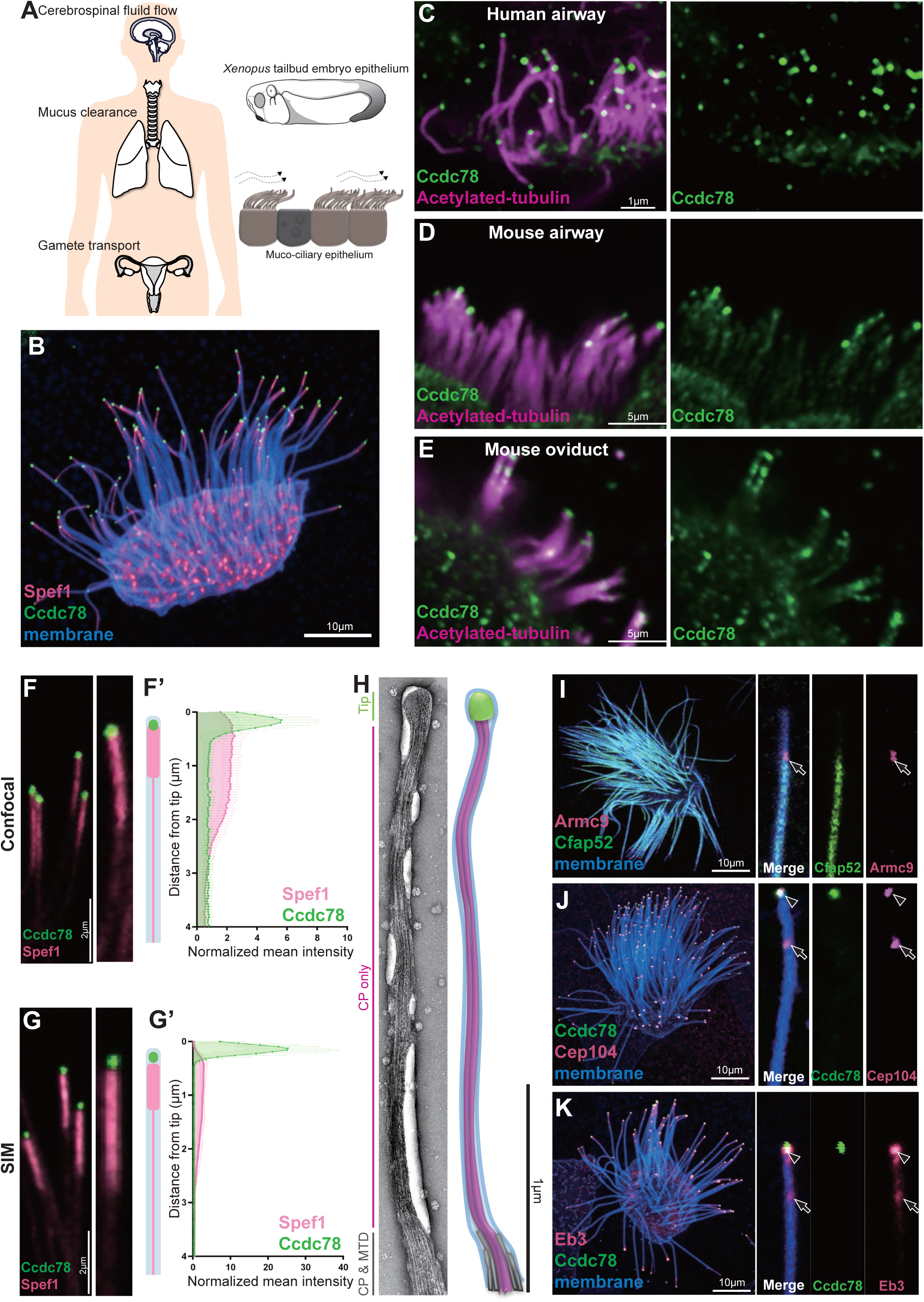
Ccdc78 localizes to the extreme distal tip of MCC motile cilia. (A) Schematic representations of the role of MCCs in the human brain, respiratory tract and reproductive system. The tailbud stage of *Xenopus* embryo is covered by muco-ciliary epithelium, which resembles mammalian MCCs. (B) Image of *Xenopus* MCC labeled with Spef1-RFP (magenta), GFP-Ccdc78 (green) and membrane-BFP (blue). Scale bar represents 10μm. (C) Immunofluorescence image of human trachea epithelial cells stained with anti-acetylated tubulin (magenta) and anti-Ccdc78 (green). Scale bar represents 1μm (D-E) Imaging of mouse airway MCCs (D) and oviduct MCCs (E) stained with anti-acetylated tubulin (magenta) and anti-Ccdc78 (green). Scale bars represent 5μm. (F) Confocal image of cilia labeled with GFP-Ccdc78 (green) and Spef1-RFP (magenta), magnified view of the cilium on the right with cartoon (F’) Quantification of fluorescent intensity along the axoneme shown in right, the mean intensity was normalized by average intensity (n=40 cilia). We quantified its localization in *Xenopus* by measuring signal intensity in the distal-most four microns of individual cilia, which allowed us to ignore confounding signals as crowded cilia lay cross one another more proximally on MCCs. (G) Structural illumination microscopy (SIM) image of cilia labeled with GFP-Ccdc78 (green) and Spef1-RFP (magenta), magnified view of the cilium on the right with cartoon. (G’) Quantification of fluorescent intensity along the axoneme shown in right (see Methods)(n=50 cilia). (H) TEM image of negatively stained *Xenopus* embryo MCC cilia with schematic model of same cilium. The tip is in green, central pair (CP) in pink and microtubule doublet (MTD) in gray. (I) Localization of mScarlet3-Armc9 (magenta, arrow) with GFP-Cfap52 (green) in a single MCC with a magnified single cilium from same cell shown at right. (J) Localization of mScarlet3-Ccdc78 (green) with GFP-Cep104 (magenta, arrowhead and arrow) in a single MCC with a magnified single cilium from same cell shown on right. (K) Localization of mScarlet3-Ccdc78 (green) with GFP-Eb3 (magenta, arrowhead and arrow) in a single MCC with a magnified single cilium from same cell shown at right.

Motile cilia in MCCs are unique for their myriad well-described specializations, from the rootlets and basal feet anchoring basal bodies to the carefully patterned positions of dynein motors along the axoneme^2^. Far less is known, however, about the specializations at the distal tips of MCC cilia^6^. This dearth of knowledge stems in part from the fact that these structures are far more variable than are other elements of motile cilia. Indeed, tip structures vary between species, between different tissues, and even between developmental stages of an organism^6^. In MCCs, the ends of all microtubules (doublets, singlets, and central pair) are each “capped” by ill-defined structures variously called caps or plugs^6–8^. The molecular nature of such caps in vertebrate MCCs is not known.

Unlike vertebrate MCC cilia, the distal tips of motile cilia in unicellular organisms have recently been resolved by CryoEM (cryogenic electron microscopy). In *Tetrahymena* for example, the distal-most region of the axoneme is occupied only by the central pair, preceded by a region that also contains A-tubule singlets, with the normal 9+2 architecture finally arising about a micron from the tip^9–11^. Electron microscopy studies reveal a similar configuration in *Chlamydomonas*^12–17^. The fine structure of the tips of motile cilia in vertebrate MCCs is less defined^7,18–20^, but studies of protein localization provide entry points for understanding.

For example, end-binding proteins that localize to MT plus ends such as Eb1 and Eb3 label the tips of both primary and motile cilia in vertebrates^21–24^. Other proteins label specifically the tips of motile cilia, for example the MT bundling protein Spef1 localizes to the central pair apparatus along the length of the axoneme^9,25^ but is dramatically enriched in the tips of *Xenopus*, mouse and human MCC cilia^24,26,27^. That said, other tip proteins described in unicellular organisms, such as Armc9 and Cep104^9–11^ have yet to be resolved clearly in vertebrate MCCs. How these proteins are recruited and arranged in distal cilia remains poorly defined.

Here, we combine confocal, structured illumination and electron microscopy to provide a quantitative description of the molecular organization of the distal tips of vertebrate MCC cilia. We find that the poorly defined proteins Ccdc78 and Ccdc33 occupy a conserved, specialized domain in the extreme distal tip of MCC cilia in *Xenopus*, mouse, and human. This region is distal to and distinct from that previously associated with other ciliary tip proteins such as Spef1. *In vivo* and *in vitro* assays reveal that Ccdc78 and Ccdc33 act in concert to bundle microtubules, suggesting they stabilize the cilia tip. Despite this shared activity, these proteins nonetheless play non-redundant roles in maintaining the normal localization of all other tip proteins and loss of Ccdc78 or Cdc33 elicits distinct defects in overall cilia length control. These, in turn, elicit distinct effects on cilia beating and resultant fluid flow. We conclude that Ccdc78 and Ccdc33 represent a novel module that organizes and stabilizes the architecture of the distal tip of motile cilia in vertebrate MCCs, thereby ensuring robust and effective cilia-mediated fluid flow.

## Results

### Ccdc78 defines a novel domain at the extreme distal tip of MCC motile cilia

The MCCs of the *Xenopus* embryo epidermis accurately reflect the biology of mammalian MCCs^28^ (Fig. 1A), and we have exploited these very large cilia as a platform for screening the localization of poorly understood proteins^27,29^. Recently, we observed that a GFP-fusion to Ccdc78 specifically marked the tips of motile cilia (Fig. 1B). This pattern of localization at the tip was consistently observed both during ciliogenesis and in mature cilia (Supp. Fig. 1A). Ccdc78 was previously implicated in the biogenesis of basal bodies in MCCs^30,31^, and as expected our GFP fusion also marked deuterosomes in the MCC cytoplasm (Supp. Fig. 1B). Localization to the cilia tip was not an artifact of the GFP fusion or of the *Xenopus* system, as immunostaining for endogenous Ccdc78 revealed localization at cilia tips in human tracheal MCCs, as well as mouse airway and oviduct MCCs (Fig. 1C, D, E; Supp. Fig. 1C).

The domain of Ccdc78 localization in cilia tips was clearly distinct from that of previously described proteins such as Spef1, which is also enriched in the distal ends of MCC cilia in humans, mice, and *Xenopus*^24,26,27^. Spef1 marks an elongate domain, while Ccdc78 was far more restricted, marking a roughly circular domain in the extreme distal tip (Fig. 1B, F). Structured illumination super-resolution microscopy indicated that the regions marked by Spef1 and Ccdc78 enrichment were largely non-overlapping (Fig. 1G). To quantify these regions, we measured pixel intensity for each marker along the distal four microns of cilia, since frequent overlap of the densely packed axonemes on MCCs made intensity measurements of more proximal regions impossible. Quantification in both confocal and structured illumination microscopy indicated that while Spef1 is enriched in the distal 2.25 microns, Ccdc78 was restricted to the most distal <0.2 microns (Fig. 1F’, G’)

In vertebrate MCCs, motile cilia are characterized by a conserved structure of nine outer doublets and two microtubules forming a central pair (9+2) that runs along the axoneme length. However, the structure of the distal tips of motile cilia varies widely between species and even cell types^6^ and the ultrastructure of the tips of *Xenopus* epidermal MCCs is unknown. Using negative stain electron microscopy (EM), we consistently observed that the extreme distal tip was characterized by a bulb-like structure occupying roughly the final 0.15 (+/- 0.02) microns of each cilium (Fig. 1H, green; Supp. Fig. 2A). This bulb was followed proximally by a narrow “neck” region roughly 2.2 microns in length (Fig. 1H, pink; Supp. Fig. 2B), after which point the axoneme quickly widens (Fig. 1H, gray).

The size and position of the distal bulb observed in EM precisely matches that of the region marked by Ccdc78, while the neck region precisely matches that of Spef1 (Fig. 1F-H). Given that Spef1 interacts specifically with central pair microtubules^9,25^, we interpret the two-micron long neck region in our EM as a central pair extension similar to that described at the distal ends of 9+2 cilia of *Tetrahymena*^9–11^. Our data thus suggest that the outer doublets of *Xenopus* MCC cilia terminate roughly 2.25 microns from the cilia tip (Fig. 1H, bottom of panel, gray).

To confirm this interpretation, we labelled *Xenopus* MCCs with Armc9, which localizes to the distal ends of the outer doublet B-tubules in *Tetrahymena*^10,11^ and Cfap52, a microtubule inner protein localized along the length of B-tubules in bovine tracheal MCCs^32^. In *Xenopus*, Armc9 labeled a small region precisely corresponding with the distal end of the Cfap52 domain, and this region fell precisely 2.2 microns from the tip (Fig.1I, arrow; quantified in Supp. Fig. 2C).

As a final test, we examined the localization of Cep104 and Eb3, each of which marks the distal ends of both central pair and doublet microtubules^22,23,33,34^. Each protein marked two regions in *Xenopus* MCCs, the extreme distal tip marked by Ccdc78 and the region 2.2 microns from the tip that is marked by Armc9 (Fig. 1J-K, arrow, arrowhead; quantified in Supp Fig. 2D-E). From these data, we propose that the Ccdc78-positive bulb at the extreme tips of MCC cilia represents a novel domain which we propose to call the extreme distal tip (EDT). The EDT is bordered proximally in *Xenopus* by a central pair extension marked by Spef1, followed by the outer doublets roughly 2.2 microns from the tip.

### Ccdc78 is required for localization of distal tip proteins in MCCs

To explore the function of Ccdc78, we performed knockdown (KD) using anti-sense morpholino-oligonucleotides that efficiently disrupted the splicing of Ccdc78 (Supp. Fig. 3A). Consistent with its known role in deuterosome function^30,31^, Ccdc78 KD significantly reduced the number of basal bodies in MCCs (Supp. Fig. 3B, C and E). This phenotype was rescued by re-expression of Ccdc78 (Supp. Fig. 3D, E), confirming the KD was specific.

Importantly, Ccdc78 KD also elicited loss of Spef1 enrichment from the distal axoneme, and this phenotype was also rescued by re-expression of Ccdc78 (Fig. 2-D). To better understand this phenotype, we examined Cep104 and Eb3 localization, finding that the normal two peaks collapsed to a single peak for each marker after Ccdc78 KD, and this peak was positioned at the very distal end of cilia (Fig. 2E-H; Supp. Fig 3F-I). Moreover, the B-tubule-end protein Armc9, normally present in a single domain two microns proximal to the tip was radically re-positioned, marking the very distal ends of cilia after Ccdc78 KD (Fig. 2I-L).

**Figure 2.**
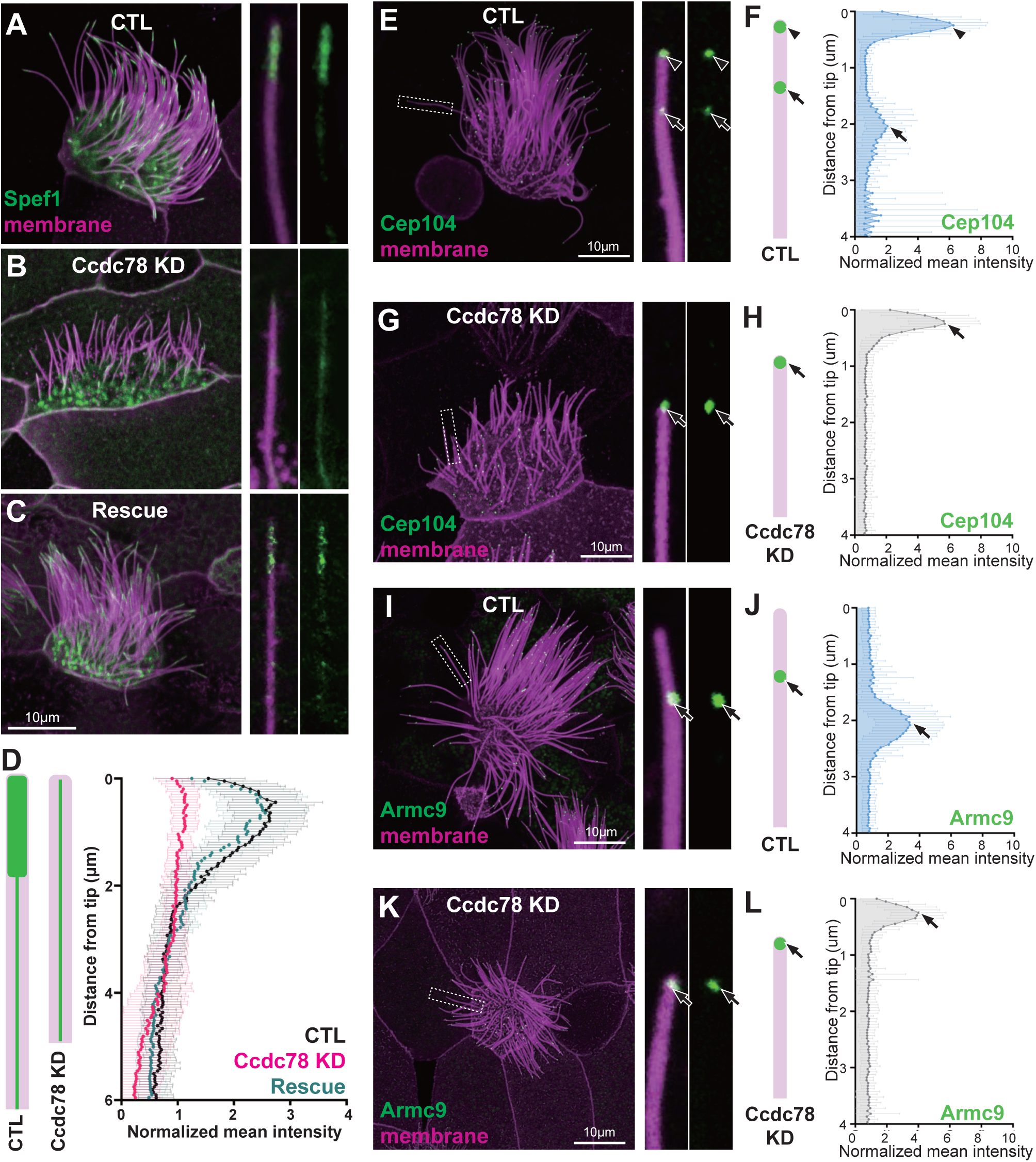
Ccdc78 is required for localization of distal tip proteins in MCC. (A-C) Image of GFP-Spef1 (green) and membrane-RFP (magenta) in *Xenopus* MCCs in control (A), Ccdc78 MO-injected (B) or rescued embryos (C) with magnified view of cilium on the right. Scale bars represent 10μm. (D) Schematic cartoon of Spef1 distribution in control and Ccdc78 MO-injected cilium. Quantification of fluorescent intensity of GFP-Spef1 along the axoneme was normalized by average intensity in control, Ccdc78 MO injected and rescued embryos. (n=58 cilia (E-H) Localization of distal proteins in control (E-F) and Ccdc78 MO injected (G-H) embryos with magnified view and schematic cartoon with quantification of normalized mean intensity on the right. Embryos were injected with Membrane-RFP (magenta) and GFP-Cep104 (green, arrowhead and arrow). Scale bars represent 10μm. (n=80 cilia (I-L) Localization of distal proteins in control (I-J) and Ccdc78 MO injected (K-L) embryos with magnified view and schematic cartoon with quantification of normalized mean intensity on the right. Embryos were injected with Membrane-RFP (magenta) and GFP-Armc9 (green, arrow). Scale bars represent 10μm. (n=80 cilia)

We interpret these results together to mean that Ccdc78 KD elicits loss of the entire distal domain that is normally marked by Spef1 and normally extends 2.2 microns beyond the Armc9 domain. Thus, we conclude that after Ccdc78 KD, the central pair extension is eliminated and that outer doublet and central pair microtubules terminate together at the very end of these defective cilia.

### Ccdc78 interacts physically and functionally with Ccdc33 in the MCC cilia tip

Ccdc78 is a coiled-coil protein with no other domains to suggest its function. To gain new insights, we searched for interacting proteins using *in vivo* affinity purification from *Xenopus* MCCs and mass spectrometry (APMS). We used an MCC-specific α-tubulin promoter^35^ to express either GFP or GFP-tagged Ccdc78 specifically in MCCs; we then isolated presumptive mucociliary epithelium, cultured it until MCCs were fully developed^36,37^, and performed APMS with anti-GFP beads (Fig. 3A). Compared to GFP alone, the most strongly enriched protein in the GFP-Ccdc78-expressing cells was the bait itself, demonstrating the efficacy of the method (Fig. 3B; Supp. Table 1).

**Figure 3.**
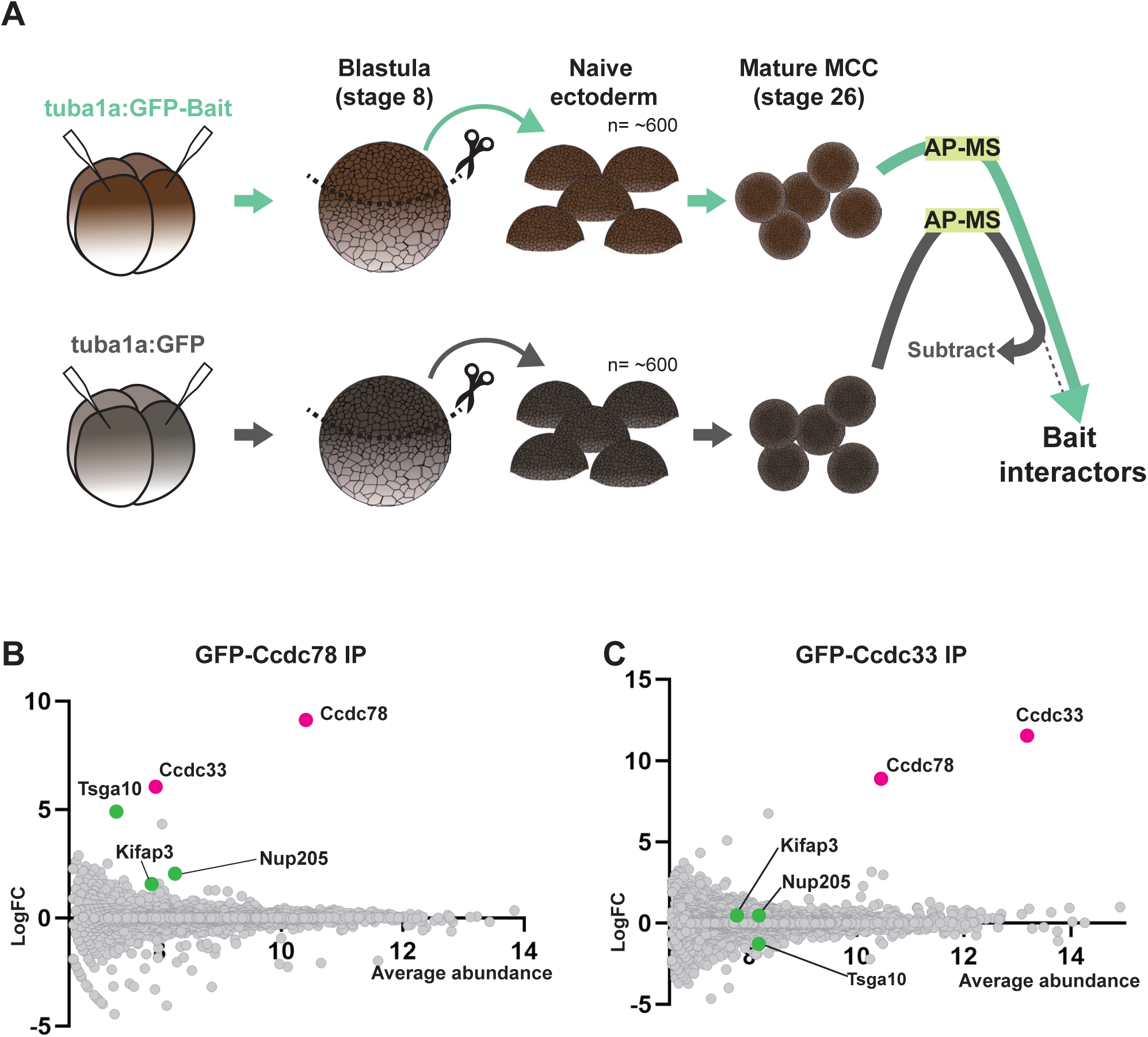
Ccdc78 interacts physically and functionally with Ccdc33 in the MCC cilia tip. (A) Workflow for AP-MS performed using *Xenopus* animal caps. (B-C) MA plot of log2 fold change and average abundance in comparison between GFP and GFP-Ccdc78 (B) or GFP-Ccdc33 (C) immunoprecipitation enriched proteins. Highly enriched proteins are labeled with magenta (Ccdc78 and Ccdc33) and green (Tsga10, Nup205 and Kifap3).

Consistent with Ccdc78’s known role in the cytoplasm^30,31^, Ccdc78 APMS enriched for several proteins acting at the base of cilia, including Nup205^38^ and Tsga10^39,40^ (Fig. 3B, green; Supp. Table 1). Other interactors were consistent with the axonemal localization for Ccdc78, such as the IFT kinesin subunit Kifap3 (Fig. 3B, green; Supp. Table 1). Our APMS did not, however, identify known vertebrate cilia tip proteins such as Spef1 or Eb3 (Supp. Table 1).

Rather, the most highly enriched prey protein was the largely uncharacterized Ccdc33 (Fig. 3B, pink). We previously identified *ccdc33* as a direct target of the ciliary transcription factor Rfx2 and found it localized to axonemes^29,41^, but another group has suggested the protein localizes to peroxisomes^42^, and nothing is known of Ccdc33 function. We therefore confirmed the interaction using reciprocal APMS with Ccdc33 as bait and found that Ccdc78 was the most strongly co-enriched prey protein (Fig. 3C, pink). Cytoplasmic Ccdc78-interactors such as Nup205 and Tsga10 were not enriched in Ccdc33 APMS (Fig. 3C, green; Supp. Table 2), consistent with Ccdc33 and Ccdc78 interacting specifically in the axoneme.

Consistent with this idea, GFP-Ccdc33 localized specifically to the EDT of cilia at all stages of ciliogenesis in *Xenopus* MCCs (Fig. 4A) and was not present the MCC cytoplasm (Supp. Fig. 4A-B). A similar result was observed for endogenous Ccdc33 in human tracheal epithelia cells differentiated in culture (Fig. 4B; Supp. Fig. 4C). Z-projection of co-stained human MCCs confirmed that endogenous Ccdc33 and Ccdc78 co-localized at cilia tips, while Ccdc78 alone was also enriched in the apical cytoplasm (Supp. Fig. 4D).

**Figure 4.**
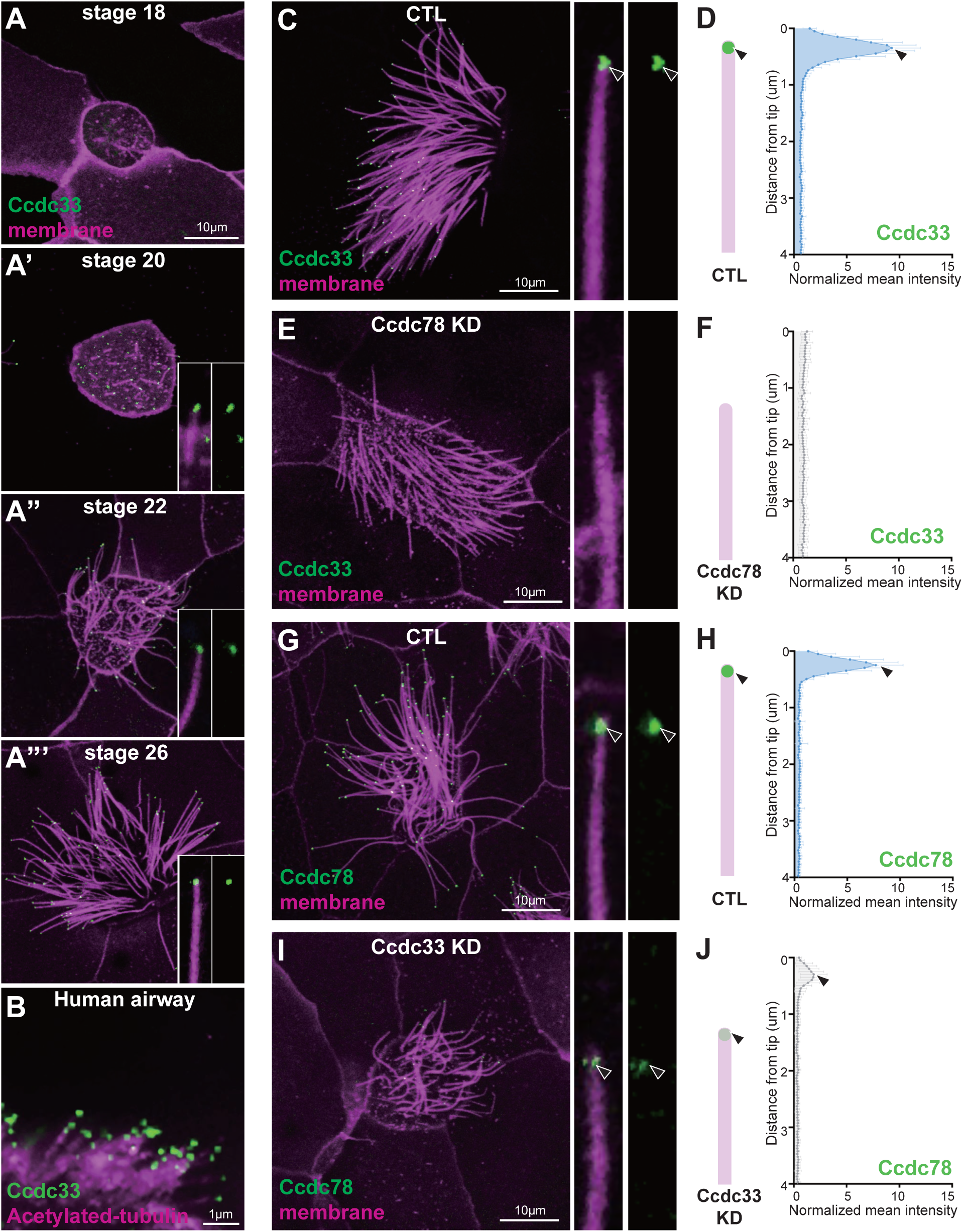
Ccdc33 localizes to the extreme distal tip of MCC cilia. (A-A”’) Localization of GFP-Ccdc33 (green) with RFP-membrane (magenta) during ciliogenesis on *Xenopus* embryo epithelium. Magnified view of cilium on the bottom right. Scale bar represents 10μm. (B) Human tracheal epithelial cell multi-cilia stained with anti-acetylated tubulin (magenta) and anti-Ccdc33 (green) antibody. Scale bar represents 1μm. (C-F) *Xenopus* MCC was injected with membrane-RFP (magenta) and GFP-Ccdc33 (green, arrowhead) in control (C-D) and Ccdc78 KD (E-F) embryo with magnified view and schematic cartoon with quantification of normalized mean intensity on the right. Scale bar represents 10μm. (n=60 cilia) (G-H) *Xenopus* MCC injected with membrane-RFP (magenta) and GFP-Ccdc78 (green, arrowhead) in control (G-H) and Ccdc33 MO-injected (I-J) embryo with magnified view and schematic cartoon with quantification of normalized mean intensity on the right. Scale bar represents 10μm. (n=60 cilia)

The co-localization of Ccdc33 and Ccdc78 at the EDT prompted us to test for functional interactions. We found that Ccdc78 KD eliminated the localization of GFP-Ccdc33 from the EDT (Fig. 4C-F). This KD did not reduce total protein levels of GFP-Ccdc33 in western blots, suggesting a specific effect on localization (Supp. Fig. 5A). Reciprocally, we developed KD reagents that disrupted splicing of Ccdc33 (Supp. Fig. 5B), and these severely disrupted but did not eliminate the EDT localization of Ccdc78 (Fig. 4G-J, Supp. Fig. 5C).

Moreover, Ccdc33 KD reduced but did not eliminate the Spef1-enriched distal domain (Fig. 5A, B and D). This effect was rescued by the re-expression of Ccdc33 mRNA (Fig. 5C, D), demonstrating the specificity of the KD. Other effects of Ccdc33 KD paralleled those of Ccdc78 KD, with loss of the distal peaks of Cep104 and Eb3, and Armc9 now abutting the cilia tip (Fig. 5E-M; Supp. Fig. 5D-G). Unlike Ccdc78, however, Ccdc33 KD had no effect on basal body numbers (Supp. Fig. 5H-J), again consistent with our proteomic and localization data suggesting Ccdc33 functions only at the cilia tip. Thus, Ccdc78 and Ccdc33 act in concert and non-redundantly to control the molecular architecture of the distal regions of vertebrate MCC cilia.

**Figure 5.**
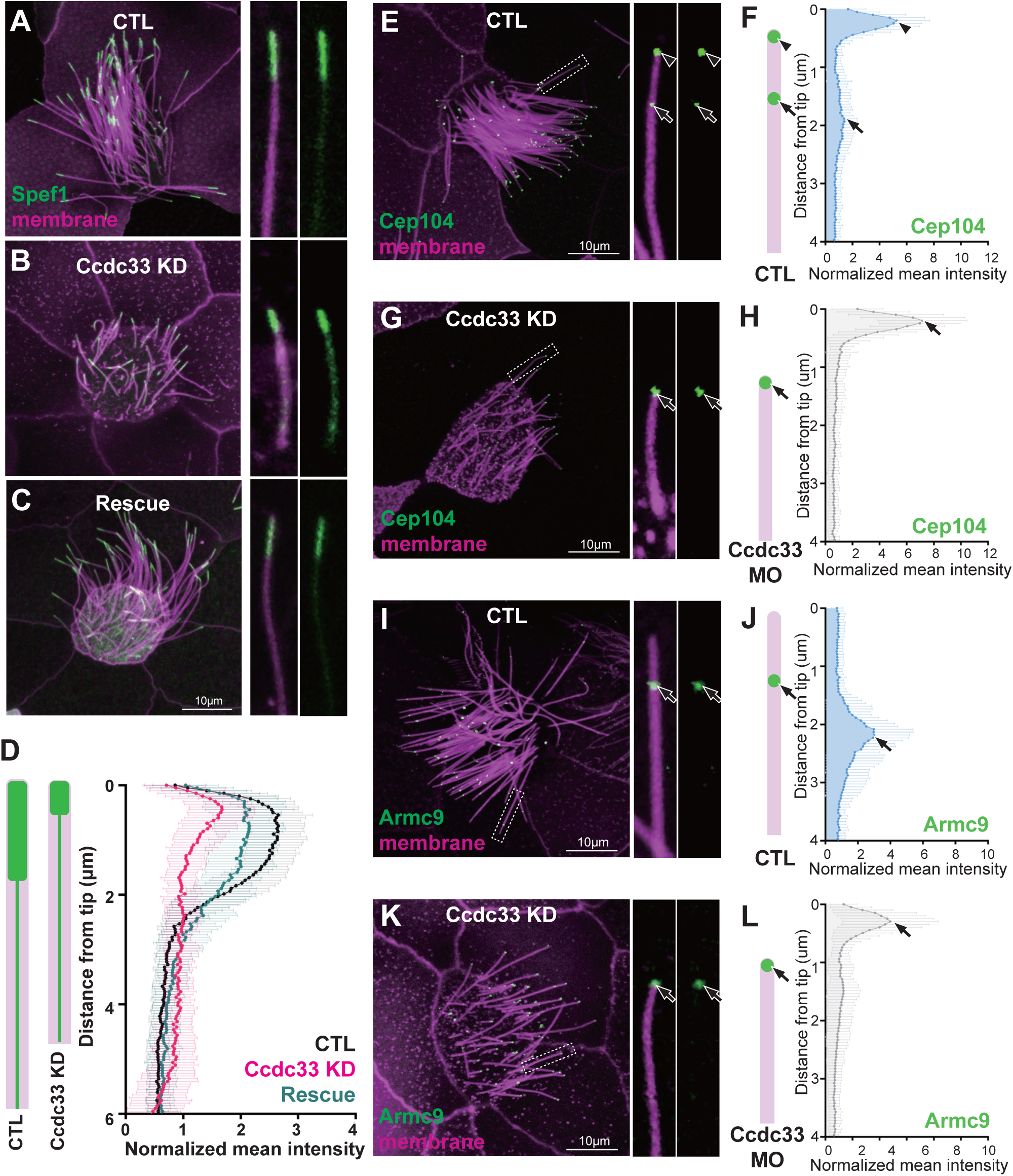
Ccdc33 is required for localization of distal tip proteins in MCC cilia. (A-C) Image of GFP-Spef1 (green) and membrane-RFP (magenta) injected *Xenopus* multi-cilia in control (A), Ccdc33 MO-injected (B) and rescued (C) embryos. Scale bars represent 10μm. (D) Schematic cartoon of Spef1 distribution in control and Ccdc33 MO-injected cilium. Quantification of fluorescent intensity of GFP-Spef1 along the axoneme normalized by average intensity in control, Ccdc33 MO-injected and rescued embryos. (n=65 cilia) (E-H) Localization of distal proteins in control (E-F) and Ccdc33 MO-injected (G-H) embryos embryo with magnified view and schematic cartoon with quantification of normalized mean intensity on the right. Embryos were injected with membrane-RFP (magenta) and GFP-Cep104 (green, arrowhead and arrow). Scale bars represent 10μm. (n=85 cilia) (I-L) Localization of distal proteins in control (I-J) and Ccdc33 MO-injected (K-L) embryo with magnified view and schematic cartoon with quantification of normalized mean intensity on the right. Embryos were injected with membrane-RFP (magenta) and GFP-Armc9 (green, arrow). Scale bars represent 10μm. (n=85 cilia)

### Ccdc78 and Ccdc33 display microtubule bundling activity *in vivo* and *in vitro*

As we considered the possible molecular functions of Ccdc78 and Ccdc33, we considered that Spef1 is known to bundle microtubules^25,43^, which in turn is thought to protect cilia tips from the high levels of mechanical strain experienced during beating^44,45^. We therefore assessed microtubule bundling by Ccdc78 or Ccdc33.

To this end, we uniformly labeled cells in the *Xenopus* epidermis with membrane-BFP and the well-defined microtubule reporter GFP-EMTB^46,47^ uniformly. We then mosaically overexpressed Ccdc78 in clones of cells marked by co-expressed memRFP (Fig. 6A)^48^. We observed that memRFP-positive clones displayed robust bundling of microtubules that was not observed in cells outside the clone (Fig. 6B; Supp. Fig. 6A and 6C-E). Mosaic expression of Ccdc33 elicited the same effect (Fig. 6C; Supp.Fig. 6B and 6C-E). We quantified local thickness of microtubule bundles inside clones expressing either protein and found them to be significantly increased compared to controls. (Fig. 6D-F). Expression of Ccdc78 or Ccdc33 did not affect the total level of GFP-EMTB protein (Supp. Fig. 6A, B) and uniform overexpression of either protein also caused marked bundling of microtubules (Supp. Fig. 6C-E).

**Figure 6.**
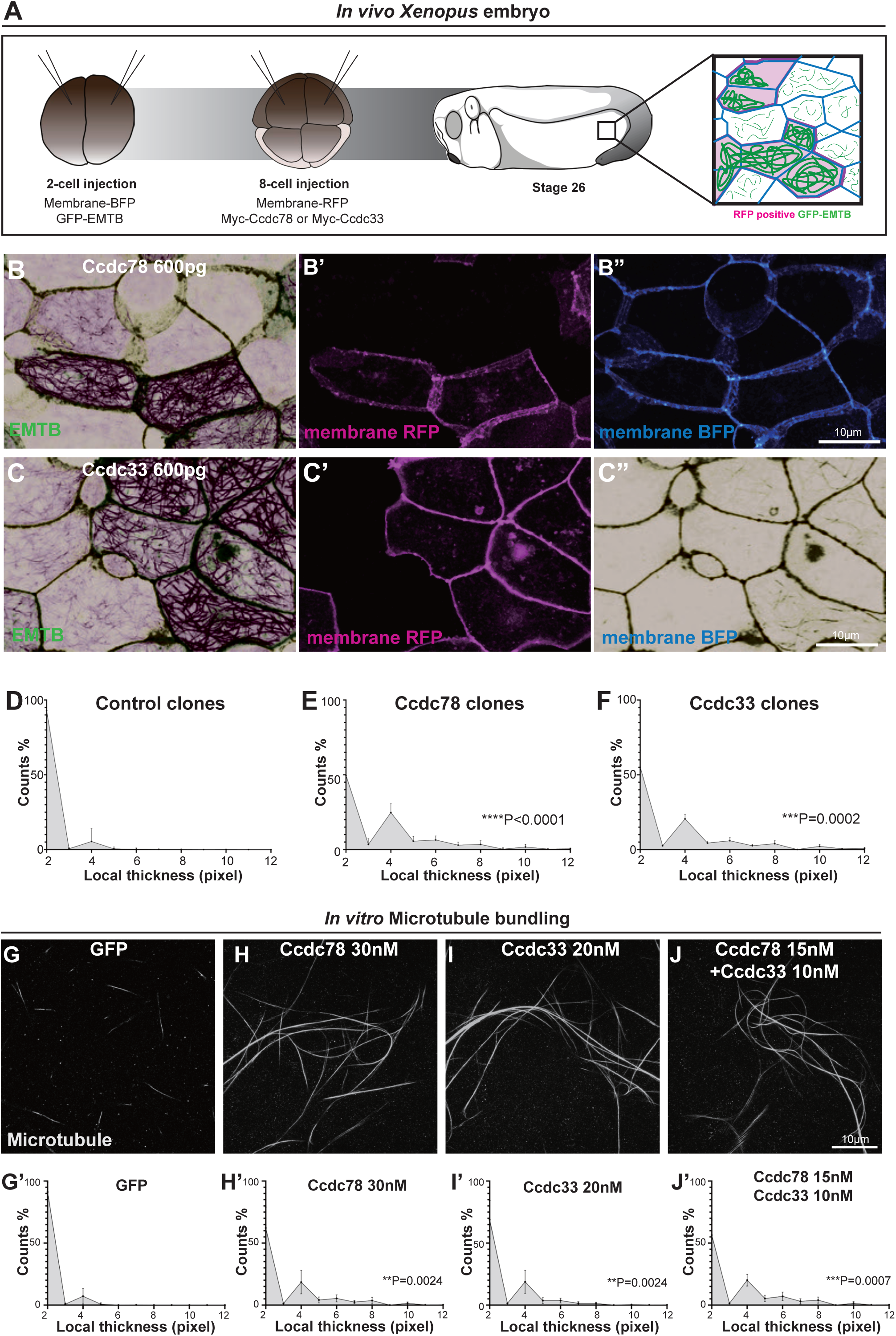
Ccdc78 and Ccdc33 display microtubule bundling activity *in vivo* and *in vitro*. (A) Workflow for generating mosaic expression in *Xenopus* embryo (B-C) Image of *Xenopus* embryo goblet cells on epithelium expressing GFP-EMTB (green), membrane-BFP (blue), and membrane-RFP(magenta) in embryos injected with Myc-Ccdc78 600pg (B-B”) and Myc-Ccdc33 600pg (C-C”). Cells expressing membrane-RFP correspond to those overexpressing Myc-ccdc78 or Myc-ccdc33, indicating mosaic expression. Scale bars represent 10μm. (D-F) Quantification of local thickness (n=26). The distribution of local thickness was compared between control (D) and Ccdc78 600pg (E) or Ccdc33 600pg (F) using Kolmogorov-Smirnov test (KS-test), the KS-distance and P-value are shown on right. CTL vs Ccdc78 600pg ****P<0.005, CTL vs Ccdc33 600pg ***P=0.0002. (G-J) Image of 647-fluorophore labeled microtubules (gray) incubated with GFP (G), GFP-Ccdc78 30nM (H), GFP-Ccdc33 20nM (I) and GFP-Ccdc78 15nM with GFP-Ccdc33 10nM (J). Scale bar represents 10μm. (G’-J’) Quantification of local thickness(n=15). The distribution of local thickness was compared between control (G’) and Ccdc78 30nM (H’) or Ccdc33 10nM (I’) or GFP-Ccdc78 15nM with GFP-Ccdc33 10nM (J’). Kolmogorov-Smirnov test (KS-test), the KS-distance and P-value are shown in right. GFP vs GFP-Ccdc78 30nM **P=0.0024, CTL vs GFP-Ccdc33 20nM **P=0.0024, GFP vs GFP-Ccdc78 15nM & GFP-Ccdc33 10nM ***P=0.0007.

We then ask if these proteins could also bundle microtubules *in vitro*. We co-incubated fluorescently labeled, polymerized, and Taxol-stabilized microtubules with *in vitro* translated GFP, GFP-Ccdc78, or GFP-Ccdc33. Microtubules incubated with GFP alone formed only sparse, small bundles (Fig. 6G). By contrast, the addition of either Ccdc78 or Ccdc33 resulted in the formation of prominent bundles throughout the sample (Fig. 6H, I). The increase in bundle thickness was significant (Fig. 6G’-I’).

Finally, we asked if the two proteins might function in concert. We found that the robust bundling achieved by 30nM Ccdc78 or 20nM Ccdc33 could also be achieved by 15nM Ccdc78 together with 10nM Ccdc33 (Fig. 6J, J’). These results demonstrate that Ccdc78 and Ccdc33 possess microtubule bundling activity *in vivo* and *in vitro* and support the idea that the two proteins work in concert to stabilize the distal tip of MCC motile cilia.

### Ccdc78 and Ccdc33 control cilia length specifically in 9+2 motile cilia

Ccdc78 and Ccdc33 bundle microtubules but also control localization of a coterie of additional proteins thought to stabilize the distal cilium, a region known to be important for ciliary length control^14,49,50^. Accordingly, we found that loss of either protein was sufficient to severely disrupt cilia length in MCCs. Ccdc78 KD severely reduced the length of MCC cilia by >8 microns (∼57% of cilium length), a phenotype that was specific since it was rescued by re-expression of Ccdc78 (Fig. 7A-D). Notably, this shortening of cilia length could not be explained merely by the loss of the distal cilium described above, as the Spef1-positive domain spans only ∼2 microns (∼15% of cilium length)

**Figure 7.**
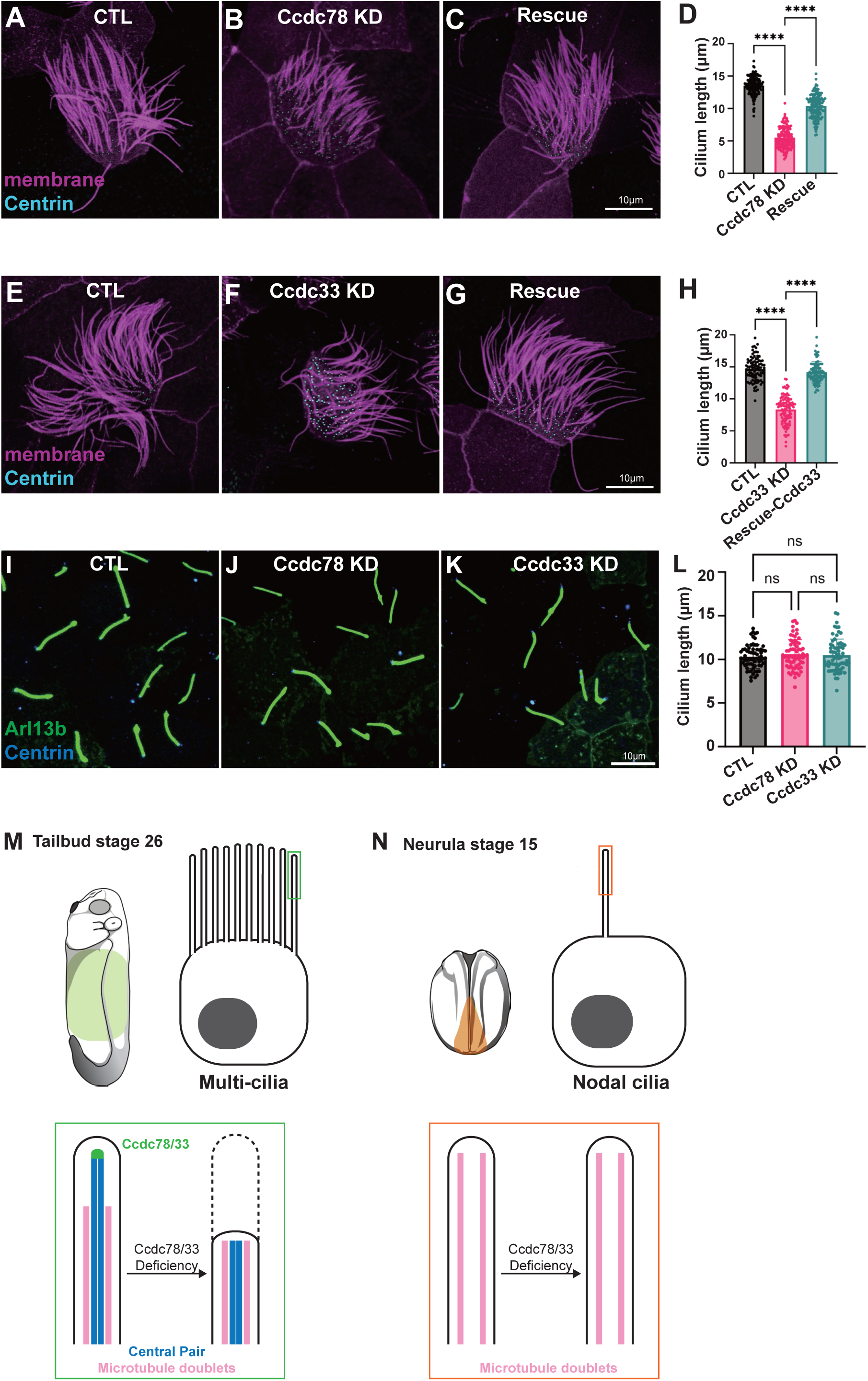
Ccdc78 and Ccdc33 are essential for length control specifically in 9+2 motile cilia. (A-C) *Xenopus* MCC injected with membrane-RFP (magenta) and Centrin-BFP (cyan) in control (A), Ccdc78 MO (B) and rescued (C) embryos. Scale bar represents 10μm. (D) Quantification of the length of cilium. ****P<0.0001. (n=147 cilia) (Ordinary one-way ANOVA) (E-G) *Xenopus* MCC injected with membrane-RFP (magenta) and Centrin-BFP (cyan) in control (E), Ccdc33 MO (F) and rescued (G) embryos. Scale bar represents 10μm. (H) Quantification of the length of cilium is shown. ****P<0.0001. (n=100 cilia) (Ordinary one-way ANOVA) (I-K) Image of *Xenopus* gastrocoel roof plate 9+0 motile cilia injected with GFP-Arl13b (green) and Centrin-BFP (blue) in control (I), Ccdc78 KD (J) and Ccdc33 KD (K) embryos. Scale bar represents 50μm. Magnified view of images are shown in right with 10μm scale bar. (L) Quantification of the length of cilia. (n=68 cilia) (Ordinary one-way ANOVA) (M-N) Schematic images of *Xenopus* embryo at tailbud stage (M) and neurula stage (N). The MCC 9+2 cilia are composed of microtubule doublets and a central pair where the extreme distal tip of central pair is labeled with Ccdc78/33. Loss of Ccdc78/33 results in disruption of distal ciliary region in MCC. On the other hand, nodal 9+0 cilia are composed of microtubule doublets only. Ccdc78/33 is not present in nodal cilia tip, thereby the 9+0 motile cilia are not affected by Ccdc78/33 deficiency.

Ccdc33 KD also had a significant -if more modest-effect on cilia length, reducing it by ∼6 microns; this effect too was specific since it was rescued by re-expression of Ccdc33 (Fig. 7E-H). Again, the loss of cilia length was too substantial to be explained only by loss of the distal Spef1 domain.

Finally, because our localization data suggested that Ccdc78/33 act as a cap on the central pair extension of 9+2 cilia, we asked if these proteins play any role in motile cilia that lack a central pair. We therefore examined 9+0 motile cilia in the *Xenopus* left/right organizer^51,52^, where Ccdc78 was present at the base of cilia but was not present at the tips as marked by Cep104 (Supp. Fig. 7A). Ccdc33 was present at neither the base nor the tip of these cilia (Supp. Fig. 7B). Moreover, KD of neither Ccdc78 nor Ccdc33 had any impact on the length of these motile 9+0 mono-cilia (Fig. 7I-L; Supp. Fig. 7C-E). Thus, Ccdc78 and Ccdc33 play non-redundant roles in assuring normal cilia length specifically in the 9+2 motile cilia of vertebrate MCCs (Fig. 7M, N).

### Ccdc78 and Ccdc33 loss elicit distinct defects in MCC cilia length control

In our imaging data, it was apparent that while loss of either Ccdc78 or Ccdc33 reduced overall cilia length, the two phenotypes were quite distinct. After Ccdc78 KD, all cilia on each MCC were reduced in length to roughly the same degree. The loss was on average ∼8 microns, with a standard deviation of 1.6 (Fig. 7D). By contrast, Ccdc33 KD resulted in MCCs with heterogeneous cilia lengths. Their average length was reduced by ∼6 microns, but the standard deviation was much higher with value of 2.2 (Fig. 7H).

We reasoned that this heterogeneous reduction in cilia length might be related to the partial loss of Ccdc78 that results from Ccdc33 KD (see Fig. 4G-J, above). Consistent with this idea, we found that for any given Ccdc33 KD cilium, the intensity of Ccdc78-GFP at the tip correlated well with the length of that cilium (Fig. 8A-D). Moreover, a similar correlation was found between the length of the partial Spef1 domain and the length of the cilium (Fig. 8E-H). This result suggests that Ccdc33 elicits its effect on cilia length in part via its ability to recruit Ccdc78 to the tips of cilia and that an intact EDT is required for normal length of the Spef1 domain and normal length of the cilium.

**Figure 8.**
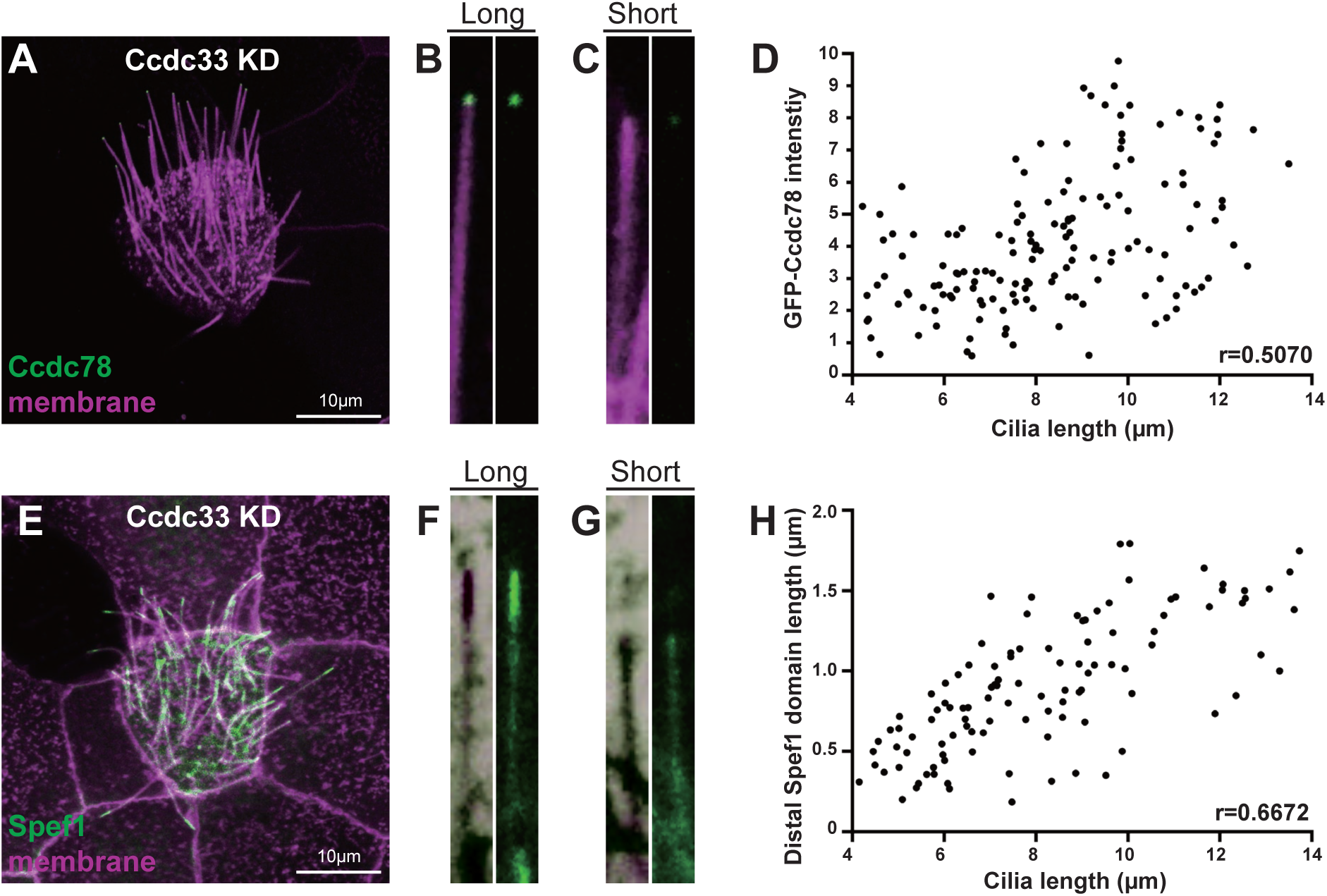
Loss of Ccdc33 elicits a heterogeneous cilia length phenotype. (A-C) Confocal image of *Xenopus* MCC injected with membrane-RFP (magenta) and GFP-Ccdc78 (green) in Ccdc33 KD embryos. The magnified view of relatively longer (B) or shorter (C) cilium are shown on the right. Scale bar represents 10μm. (D) Correlation between cilia length and GFP-Ccdc78 intensity at the ciliary tip in Ccdc33 KD embryos MCCs. Each dot represents a single cilium. A moderate positive correlation was observed (Pearson’s correlation coefficient, r = 0.5070). (E-G) Confocal image of *Xenopus* MCC injected with membrane-BFP (magenta) and Spef1-RFP (green) in Ccdc33 KD embryos. The magnified view of relatively longer (F) or shorter (G) cilium are shown on the right. Scale bar represents 10μm. (n=150 cilia) (H) Correlation between cilia length and Spef1-RFP intensity at the ciliary tip in Ccdc33 KD embryos MCCs. Each dot represents a single cilium. A moderate positive correlation was observed (Pearson’s correlation coefficient, r = 0.6672). (n=123 cilia)

### Ccdc78 and Ccdc33 loss elicit distinct defects in cilia beating and fluid flow

The precise role of the cilia tip in cilia beating remains largely unresolved, as disruption of other cilia tip proteins is confounded by their possible roles outside the tip. Spef1, for example, is dramatically enriched distally but is also present along the entire length of the axoneme and Spef1 mutant cilia entirely lack the central pair^9,25^. Likewise, Eb1 and Eb3 bind the minus ends of even cytoplasmic microtubules^21,22^, and Cep104 is present at basal bodies as well as cilia tips^13^. Ccdc78 and especially Ccdc33 therefore provide novel entry points for examining the functional consequence of disrupting the cilia tip. Notably, the two knockdowns elicited strikingly different beating phenotypes.

In spinning disk confocal microscopy, the metachronal wave of beating cilia was expectedly normal in control MCCs. To our surprise, metachrony also appeared normal after Ccdc78 KD (Fig. 9A, B; Movie 1, 2). While the uniform reduction of cilia length was apparent in our videos, demonstrating that knockdown was effective in the samples, we nonetheless observed no qualitative defect in their beating (Fig. 9B, Movie 2).

**Figure 9.**
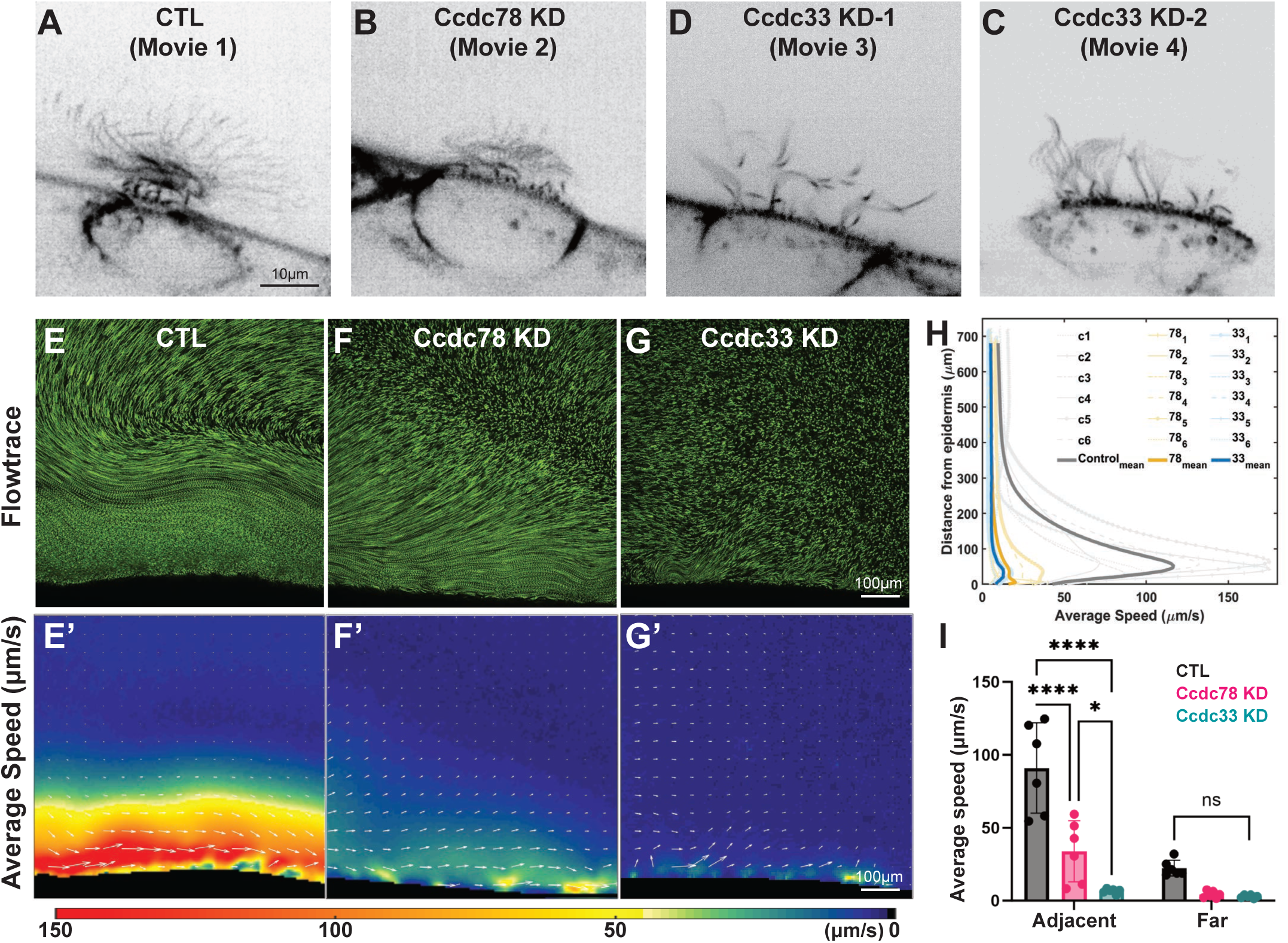
Loss of Ccdc78 and Ccdc33 elicit distinct defects in cilia beating and fluid flow. (A-D) spinning disk microscopy image of ciliary beating in control (A), Ccdc78 KD (B), Ccdc33 KD-1(C) Ccdc33 KD-2(D) MCC. Scale bar represents 10μm. (E-G) Flowtrace visualizations of one representative sample of control (E), Ccdc78 KD (F) and Ccdc33 KD (G) with 10 seconds time averaging. Scale bar represents 100μm. (E’-G’) Speed colormap corresponding to panels 9E-G overlaid with velocity vectors (white arrows). The color scale of average speed is shown at the bottom. Scale bar represents 100μm. (H) Speed profile graph showing average speed of fluid flow plotted as a function of distance from epidermis for all samples, with the mean values highlighted. (n=6 embryos) (I) Average flow speed comparison of adjacent and far regions for all samples of control, Ccdc78 KD and Ccdc33 KD. The bar refers to mean value and the error bars refer to standard errors.

By contrast, beating was visibly defective after Ccdc33 KD. In some cells, cilia appeared stiffer and metachrony was entirely absent (Fig. 9C, Movie 3). In other cells, metachrony was observed in patches where cilia length was more uniform but was absent from other regions of the same cell (Fig. 9D, Movie 4). These distinct beating patterns prompted us to assess fluid flow, the functional output of MCC cilia beating.

To this end, we imaged fluorescent tracer bead movement across the *Xenopus* embryo media using spinning disk confocal microscopy. We visualized flow using Flowtrace^53^ and quantified the data using Particle Imaging Velocimetry^54^. This analysis revealed the expected highly aligned fluid flow near the embryo surface with less robust but still obvious flow in regions farther from the surface (Fig. 9E, E’, Movie 5). Consistent with the relatively normal metachronal beating, Flowtrace revealed that flow was aligned and coordinated near surface after Ccdc78 KD, though flow speed was strongly reduced and the coordination of flow decayed quickly with distance from the surface (Fig. 9F, F’, Movie 6). By contrast, we observed a near-total absence of coordinated flow in Ccdc33 KD embryo (Fig. 9G, G’, Movie 7). These patterns were highly consistent across samples (Fig. 9H)

To quantify this effect, we calculated flow in two regions separated at the point of maximum decline in speed (Supp. Fig. 8A-D). Within the region near the surface, Ccdc78 KD reduced flow speed by roughly 50%, while Ccdc33 KD reduced speed by ∼90% (Fig. 9I, adjacent). In the region farther from the surface, the modest flow observed for controls was effectively eliminated by loss of either protein (Fig. 9I, far). These data link disruption of the EDT and the overall organization of the cilia tip to defects in cilia beating and cilia-mediated fluid flow.

## Discussion

We have defined a domain in the extreme distal tip of vertebrate MCC cilia that is defined by the presence of Ccdc78 and Ccdc33 (Fig. 1). These proteins function non-redundantly and are required for normal localization of several additional ciliary tip proteins (Fig. 2-5). Ccdc78 and Ccdc33 display robust microtubule bundling activity both *in vivo* and *in vitro* (Fig. 6). Moreover, both proteins – and by extension the EDT itself – are necessary for maintaining normal ciliary length control in MCCs (Fig. 7, 8). Finally, both are likewise required for cilia beating for MCC’s ability to generate normal fluid flows across the epithelium (Fig. 9). These data provide a new depth of understanding of the molecular architecture of MCC cilia tips and should be significant on several levels.

First, our data allow us to propose a model whereby Ccdc78 and Ccdc33 non-redundantly stabilize microtubules via their bundling activity. Since Ccdc78 is entirely necessary for Ccdc33 localization to the ciliary tip, we suggest that after Ccdc78 KD, the loss of both proteins from the tip leads to the observed uniform, severe length defect. By contrast, Ccdc33 KD elicits only a partial loss of Ccdc78 localization to the tip, which leads to the heterogeneous reduction of cilia length. And the heterogeneity of cilia length may result in more severe cilia length-dependent phenotypes.

Neither length phenotype can be explained simply by loss of the central pair extension (i.e. that marked by Spef1), so our data suggest two possibilities. First, axonemes may be destabilized in the absence of microtubule bundling by Ccdc78/33 and shortened by disassembly as a consequence of the forces associated with ciliary beating. Indeed, the whip-like movement of cilia increases mixing and velocity of the surrounding fluid most strongly near ciliary tips^44,45^, suggesting the distal axoneme is subjected to higher mechanical stress. In this light, it is notable that like Ccdc78/33, Spef1 also displays robust MT bundling and stabilizing activity *in vivo* and *in vitro*^25,43,55^. Alternatively, since elongation of the axoneme is controlled at the tip, Ccdc78/33 may somehow control the elongation machinery directly, for example via IFT turnaround^56–58^. Consistent with this notion, we identify the IFT kinesin subunit Kif3ap as a Ccdc78 interactor.

In addition, we found that Ccdc78 and Ccdc33 are present at the distal ends of *Xenopus* as well as mammalian tracheal and oviduct MCCs. This evolutionary conservation is likely very ancient because vertebrate Ccdc33 comprises coiled-coil regions and lipid-binding C2 domains, suggesting it is a direct orthologue of the C2D1 protein reported with a similar structure at the distal ends of *Tetrahymena* cilia^9,10^ (Supp. Fig. 9A-C). Likewise, a protein with homology to Ccdc78 has also been found at the distal tip of *Tetrahymena* cilia (Uniprot: I7LVY1)^10^, though that protein is characterized by a kinesin motor domain that is absent from vertebrate Ccdc78 (Supp. Fig. 9D-F). Our data also reveal that Spef1, Cep104, and Armc9 in the distal tips of *Xenopus* MCC cilia all display similar patterns to that observed in *Tetrahymena*^9–11^.

These similar patterns of protein localization are of interest because they suggest that the molecular architecture underlying 9+2 distal tips is deeply conserved, despite the fact that the ultra-structures of these regions are highly variable across species, across cell type, and even across developmental stage for a single cell type. Indeed, older EM studies suggest that a “cap” structure connects microtubules at the tip to the ciliary membrane in a wide range of vertebrate MCCs^7,17^. Since Ccdc33 contains lipid-binding C2 domains^59^, Ccdc33 and Ccdc78 may be essential components of these cap structures.

Finally, our data provide new insights into the relationships between the tip, cilia length, metachrony, and fluid flow. We find that the uniform decrease in length of cilia after Ccdc78 loss allows normal metachronal beating, but results in reduced overall fluid flow speeds (Fig. 9). This is consistent with theory suggesting that shorter cilia produce less flow^60^. More interesting are the beating defects after Ccdc33 KD. It is possible these stem directly from Ccdc33 function at the tip, but this seems unlikely because Ccdc33 is lost from the tip yet beating and metachrony are largely normal after Ccdc78 KD (Fig. 4). Another possibility is that the reduced cilia length coupled to length heterogeneity disrupts metachrony and thus severely disrupts flow even adjacent to the epithelium (Fig. 9). This interpretation is reasonable, as metachrony arises from hydrodynamic coupling^61–63^ and in theoretical models is sensitive to noise in the system^64,65^.

In light of their roles in cilia beating, it is notable that neither *CCDC78* nor *CCDC33* has previously been associated with motile ciliopathy. Rather, *CCDC78* is implicated in myopathy^66^. Not all studies support this association however,^67^ and both bulk and single-cell RNAseq in the Human Protein Atlas demonstrate that human *CCDC78* is strongly expressed in tissues that contain MCCs and at only very low levels in all other tissues, including muscle^68,69^. On the other hand, one report implicates *CCDC78* in myopathy associated with respiratory insufficiency^70^, which may be consistent with the modest defect in cilia beating. If *CCDC78* variants do contribute to myopathy, the effect may relate to the protein’s association with centrioles in MCCs, as CCDC78 was found to localize to centrosomes in HeLa cells^71^. *CCDC33* is not yet associated with any disease, but it is enriched only in tissues that contain MCCs and is co-expressed with *CCDC78* in an MCC-related cluster in scRNAseq data^68,69^.

Given that many cases of human ciliary dyskinesia go undiagnosed and that many, too, are associated with grossly normal ultrastructure of the beating machinery such as dynein arms, radial spokes, or the central pair^72,73^, our data suggest that EDT defects may explain failure of cilia beating in such patients. As such, *CCDC78* and especially *CCDC33* represent interesting new candidate loci for human motile ciliopathy.

## Figure legends

**Supplementary figure 1.**
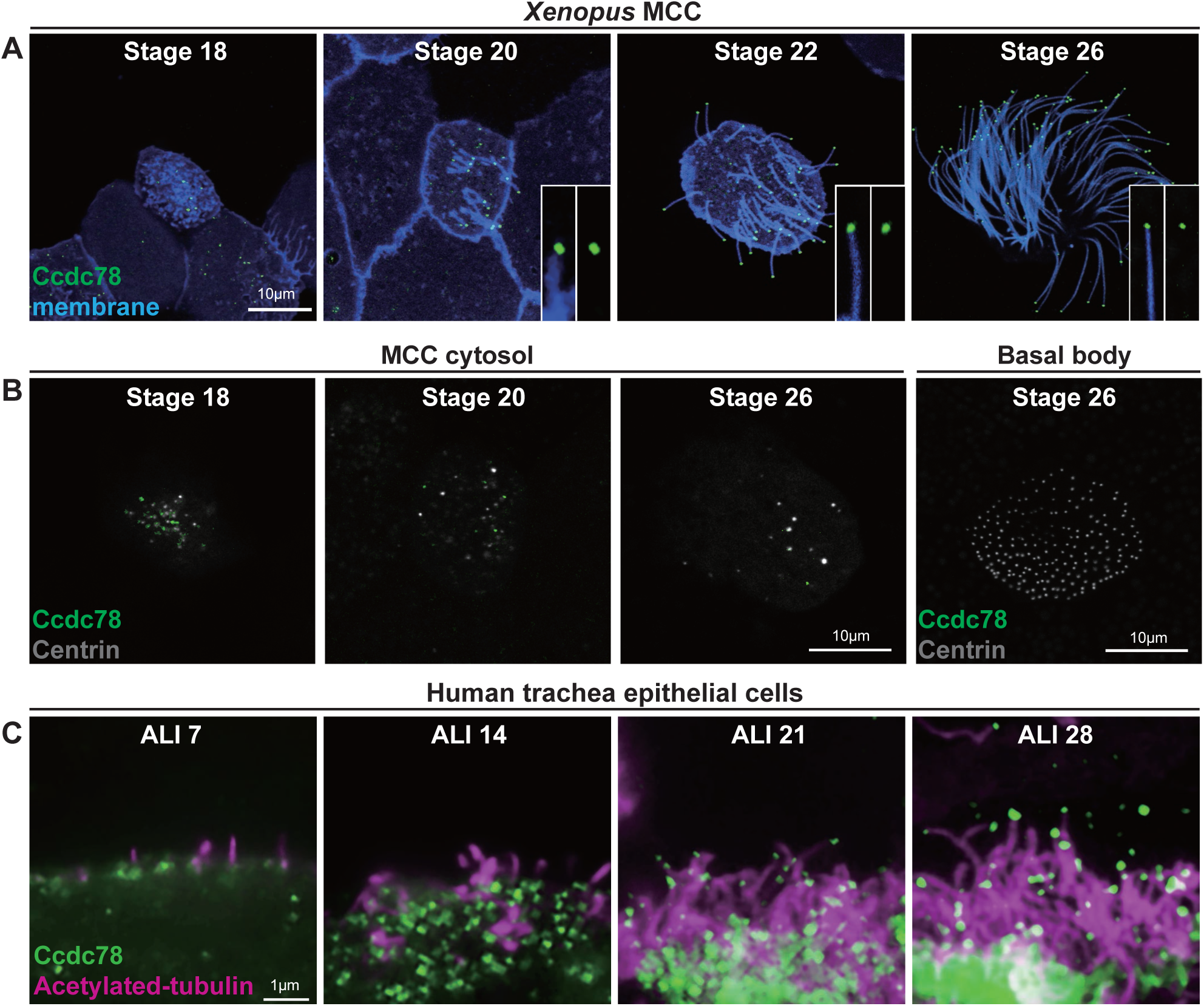
Localization of Ccdc78 in MCC cytosol during ciliogenesis. (A) Apical surface of *Xenopus* embryo epithelium of stage 18, 20, 22, and 26, showing the localization of GFP-Ccdc78 (green) with Membrane-RFP (blue). Insets show magnified views of cilia. Scale bars represent 10μm. (B) Image of *Xenopus* MCC cytosol expressed with GFP-Ccdc78 (green) and Centrin-BFP (gray) in stage 18, 20 and 26 embryos. Localization of GFP-Ccdc78 on the basal body of MCC is shown on the right. Scale bars represent 10μm. (C) Human trachea epithelial cells (HTEC) MCC stained with anti-acetylated tubulin (magenta) and anti-Ccdc78 (green) antibody during the time of ALI-culture day 7, 14, 21 and 28. Scale bar represents 1μm.

**Supplementary figure 2.**
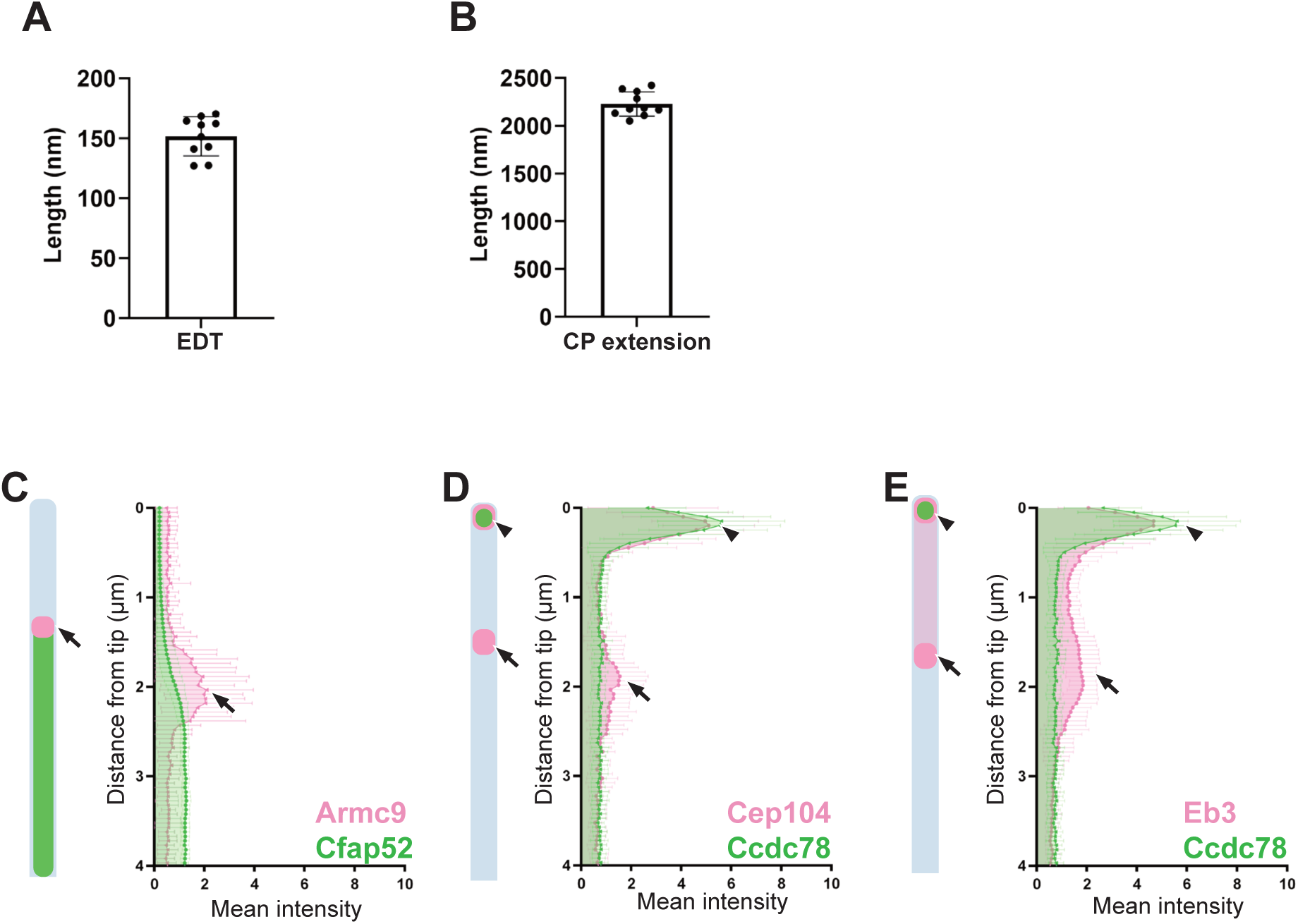
Quantification of distal ciliary proteins localization along the MCC cilia. (A) Quantification of the length of extreme distal tip (EDT) region in TEM images of *Xenopus* MCC cilia. (n=10 cilia) (B) Quantification of the length of central pair (CP) extension region in TEM images of *Xenopus* MCC cilia. (n=10 cilia (C-E) Quantification of the normalized mean intensity distribution of distal proteins Armc9 with Cfap52 (C), Cep104 (D) and Eb3 (E) shown in Fig. 1I, J, K. Arrowhead and arrow pointing at the enrichment of proteins respectively. (n=40 cilia)

**Supplementary figure 3.**
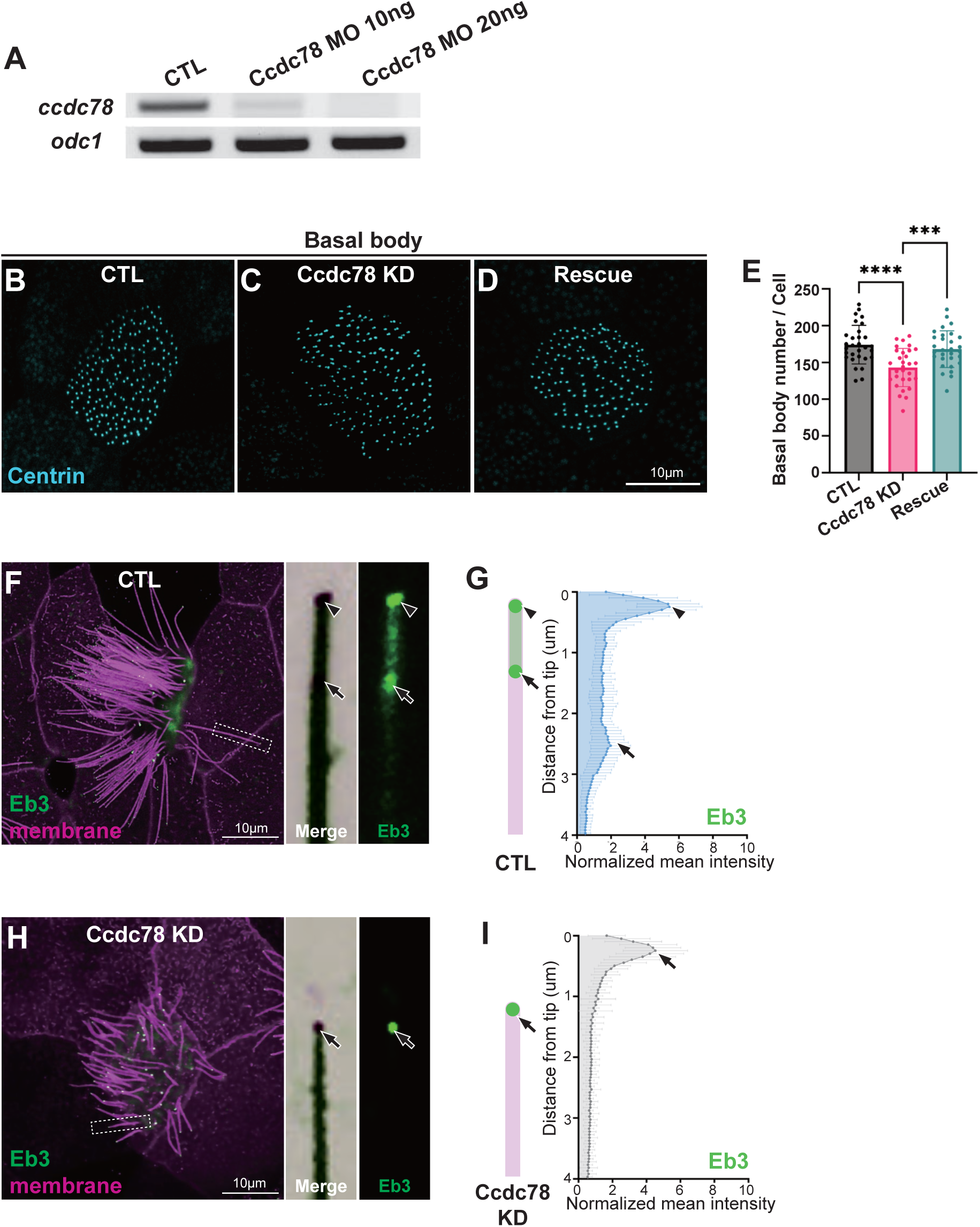
RT-PCR of Ccdc78 splice-blocking morpholino-injected embryos and quantification of distal protein distributions. (A) Gel image of RT-PCR showing *ccdc78* and *odc1* mRNA levels in control embryos and embryos injected with 10 ng or 20ng of Ccdc78 MO (B-E) Image of basal bodies expressed with Centrin-BFP (cyan) in control (B), Ccdc78 KD (C) and rescue (D) embryos. Scale bar represents 10 μm. (E) Quantification of the number of basal bodies per cell is shown on the right. ****P<0.0001, **P<0.01. (n=31 cells) (Ordinary one-way ANOVA). (F-I) Confocal image of membrane-RFP with GFP-Eb3 (arrowhead and arrow) in control (F-G) and Ccdc78 KD (H-I) MCC cilia with magnified view of cilium on the right. Quantification of the normalized mean intensity is on the right. (n=80 cilia)

**Supplementary figure 4.**
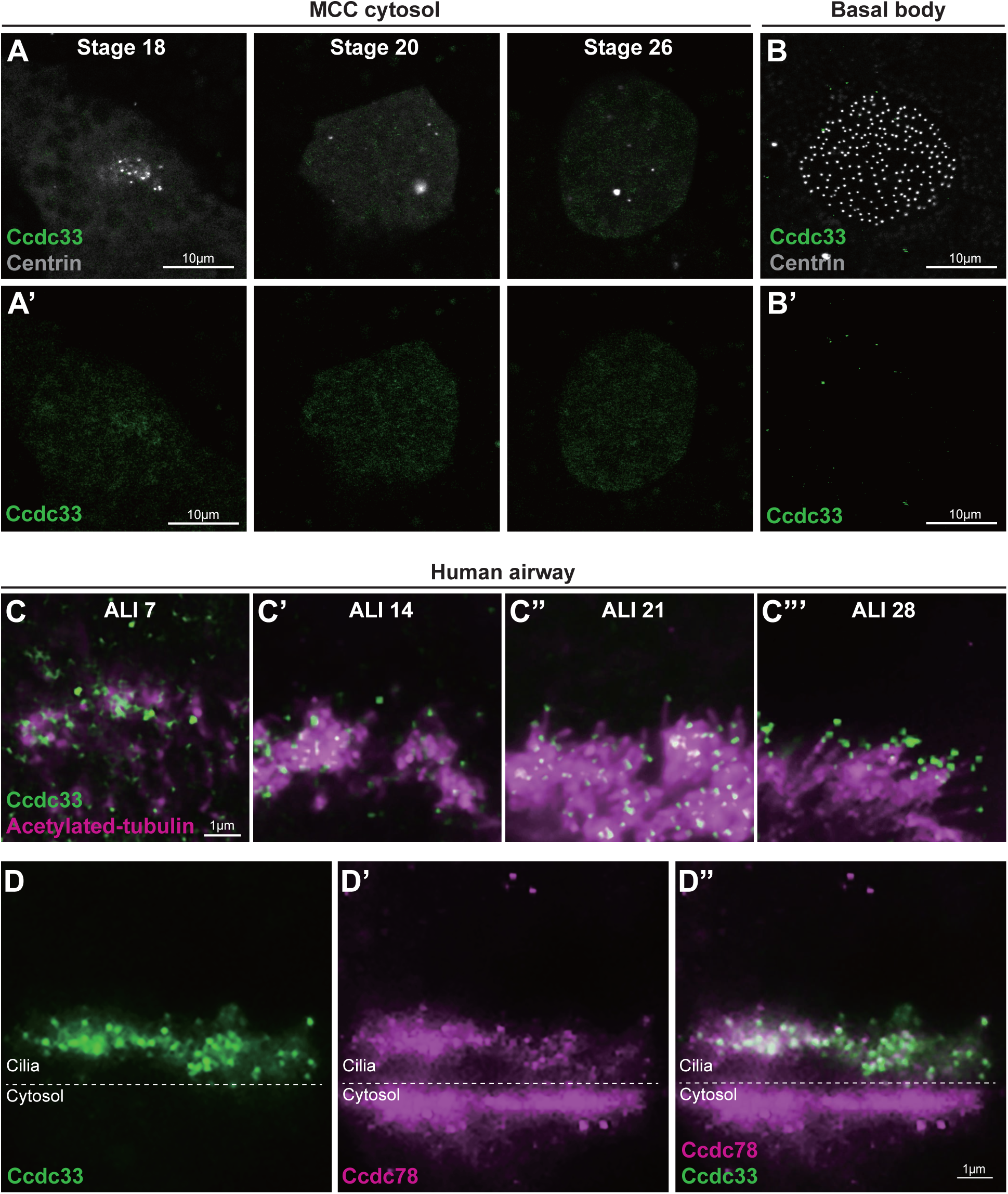
Localization of Ccdc33 in *Xenopus* MCC cytosol and HTEC. (A-B) Images of *Xenopus* MCC cytosol expressed with GFP-Ccdc33 (green) and Centrin-BFP (gray) in stage 18, 20 and 26 embryos (A). The localization of GFP-Ccdc33 on the basal body of MCC is shown on the right (B). Scale bars represent 10μm. (C) Cultured human airway MCC stained with anti-acetylated tubulin (magenta) and anti-Ccdc33 (green) antibodies during the time of ALI-culture day 7 (C), 14 (C’), 21 (C”) and 28 (C”’). Scale bar represents 1μm. (D) Cultured human airway MCC stained with anti-Ccdc78 (magenta) and anti-Ccdc33 (green) antibodies. Scale bar represents 1μm.

**Supplementary figure 5.**
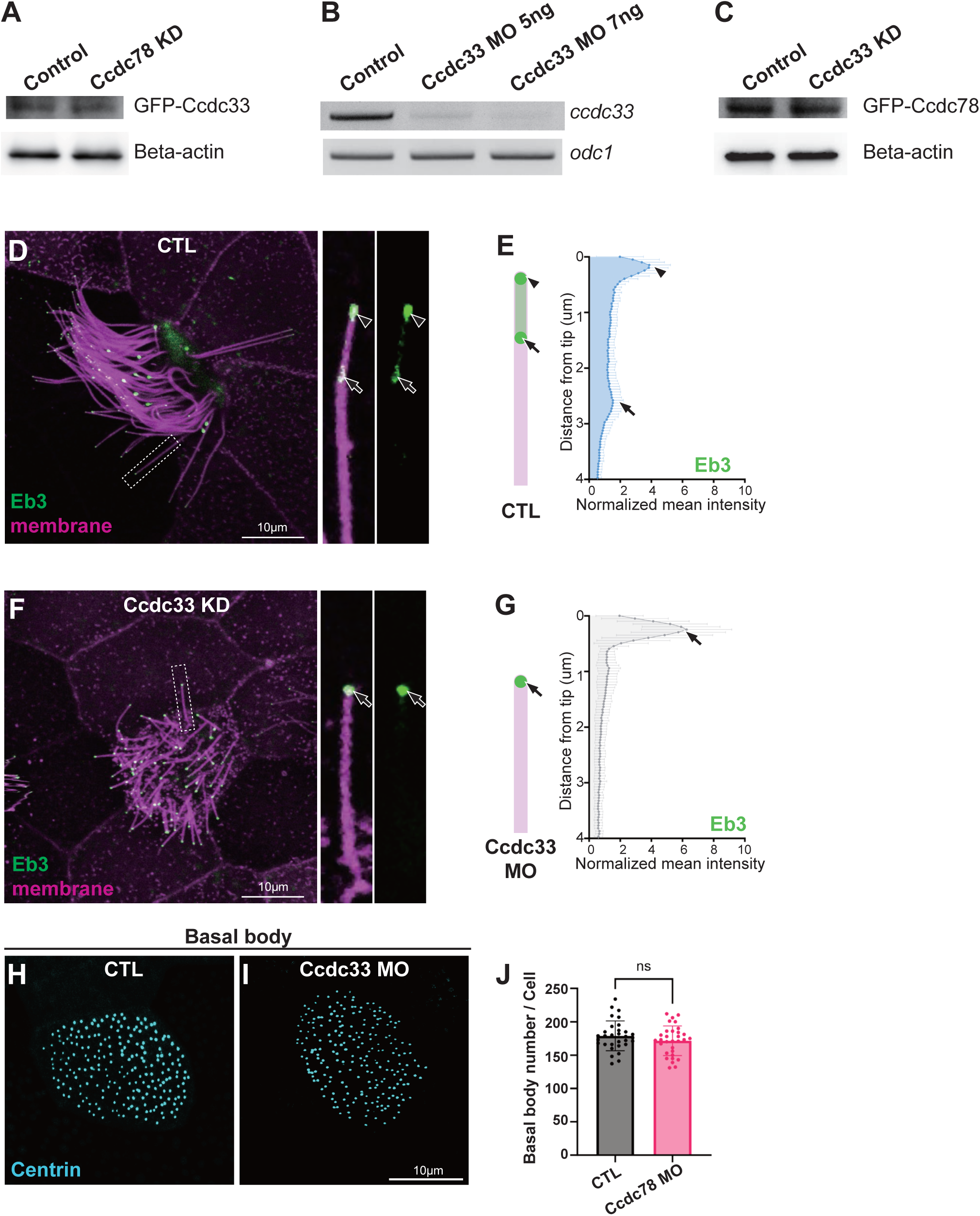
Validation of Ccdc33 splice-blocking morpholino efficiency by RT-PCR and distribution of Ccdc78/Ccdc33 intensity along the cilia after Ccdc33/Ccdc78 MO injection. (A) The total amount of GFP-Ccdc33 proteins after Ccdc78 KD was confirmed by western blot using anti-GFP and anti-beta-actin as a loading control. (B) Gel image of RT-PCR of *ccdc33* and *odc1* mRNA levels in control, Ccdc33 MO 5ng and 7ng injected embryos. (C) The total amount of GFP-Ccdc78 proteins after Ccdc33 KD was confirmed by western blot using anti-GFP and anti-beta-actin as a loading control (D-G) Confocal image of membrane-RFP with GFP-Eb3 (arrowhead and arrow) in control (D-E) and Ccdc33 KD (F-G) MCC cilia with magnified view of cilium on the right. Quantification of the normalized mean intensity is on the right. (n=80 cilia (H-I) Image of basal bodies expressed with Centrin-BFP (cyan) in control (H), Ccdc33 KD (I) embryos. Scale bar represents 10μm. (J) Quantification of the number of basal bodies per cell is shown on the right. P=0.2066. (n=30 cells) (Unpaired t-test).

**Supplementary figure 6.**
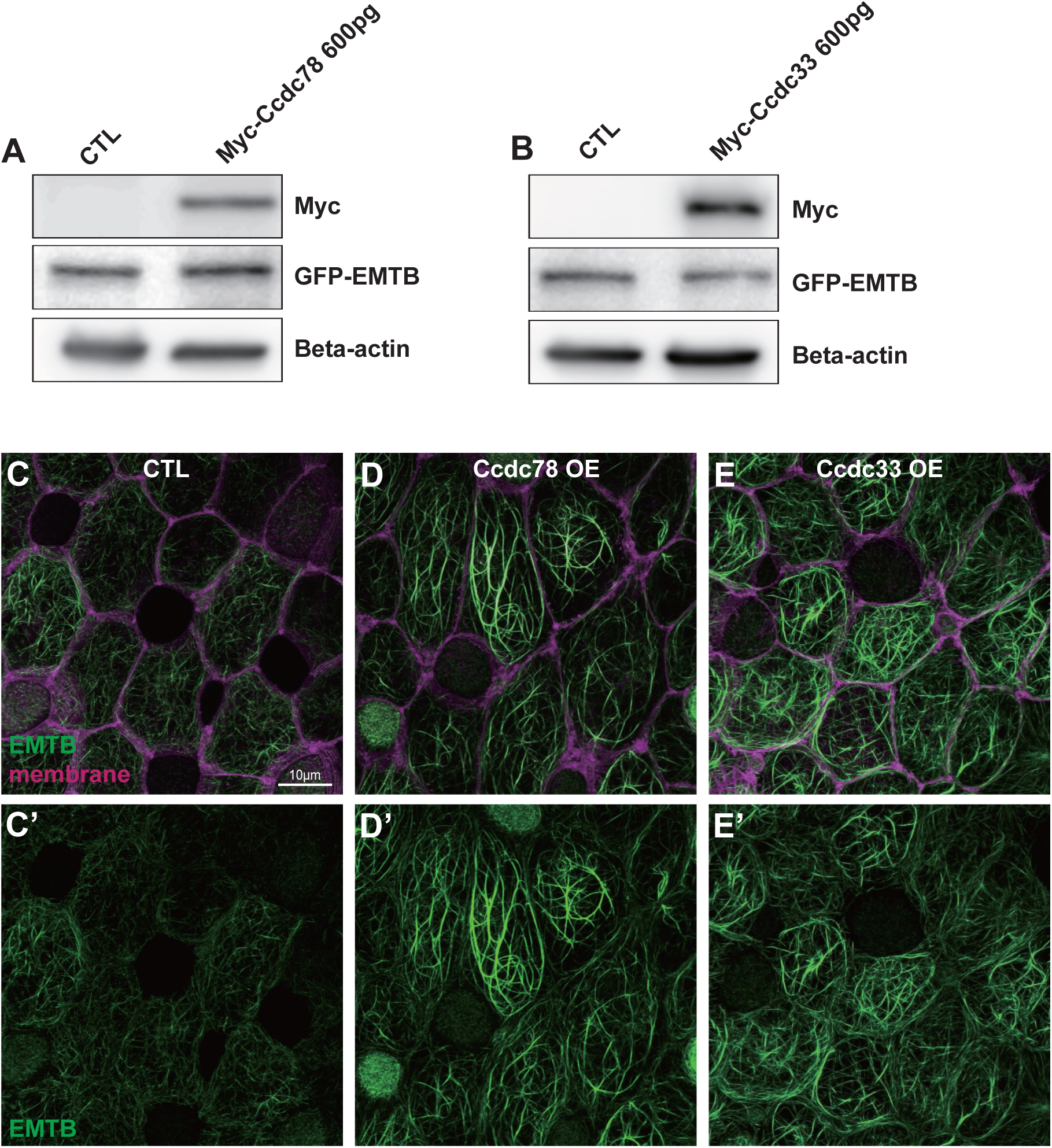
Mosaics on *Xenopus* embryo epithelium and Western blots of Myc-Ccdc78 and Myc-Ccdc33. (A-B) Western blots to validate ectopic expression of Myc-Ccdc78 (A), Myc-Ccdc33 (B), and GFP-EMTB with Beta-actin used as a loading control. (C-E) Image of *Xenopus* embryo goblet cells on epithelium expressed with GFP-EMTB (green), membrane-RFP (magenta) and overexpression (600pg) of Myc-Ccdc78 (D) or Myc-Ccdc33 (E). Scale bar represents 10μm.

**Supplementary figure 7.**
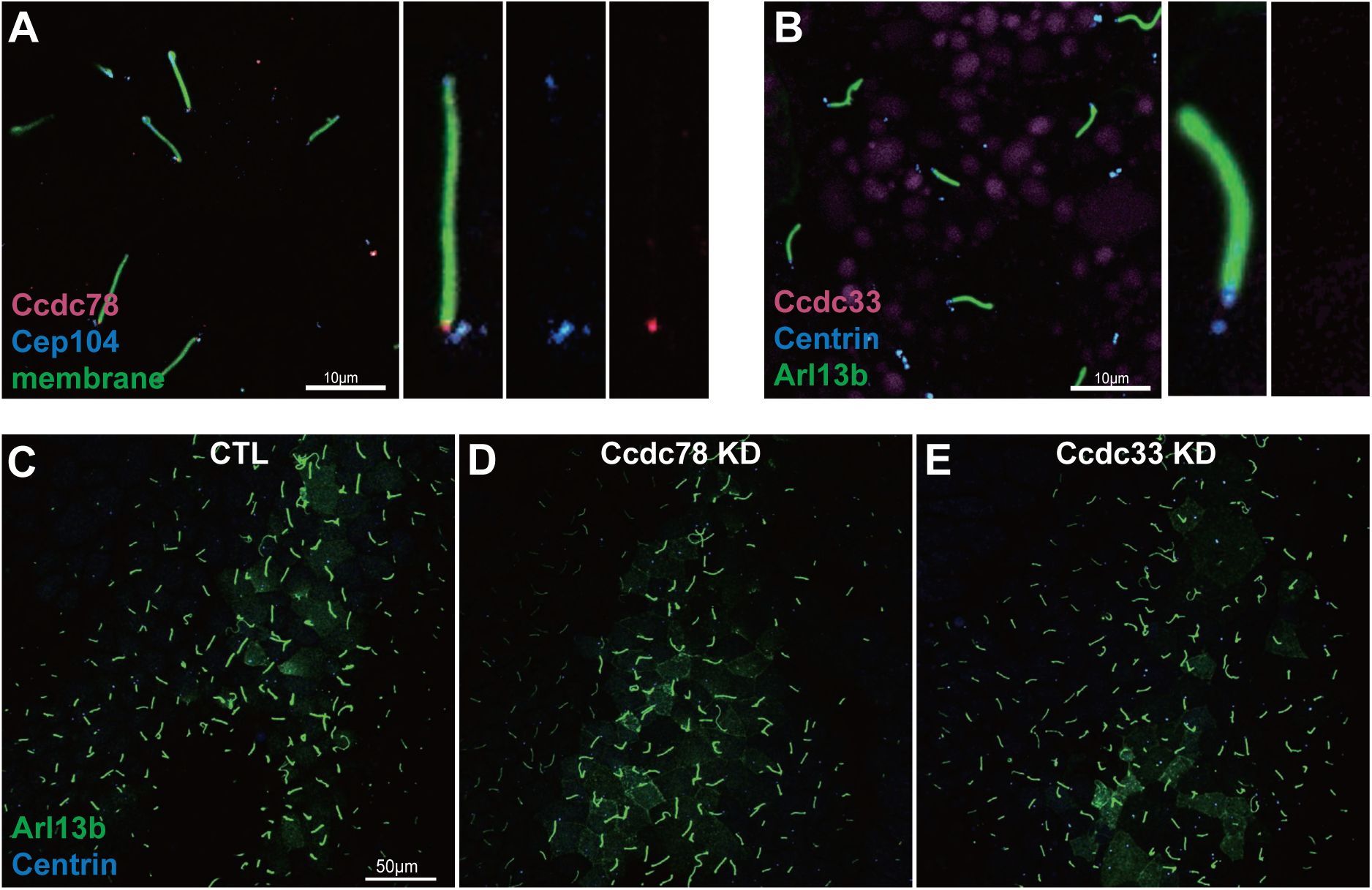
Ccdc78 and Ccdc33 specifically affect 9+2 motile cilia in MCC. (A) Localization image of mScarlet3-Ccdc78 (magenta), GFP-Cep104 (blue) and membrane-BFP (green) in 9+0 motile cilia of *Xenopus* gastrocoel roof plate. Magnified view of cilium is inserted in bottom right. Scale bar represents 10μm. (B) Localization image of mScarlet3-Ccdc33 (magenta), GFP-Arl13b (green) and Centrin-BFP (blue) in 9+0 motile cilia. Magnified view of cilium is inserted in bottom right. Scale bar represents 10μm (C-E) Image of *Xenopus* gastrocoel roof plate 9+0 motile cilia injected with GFP-Arl13b (green) and Centrin-BFP (blue) in control (C), Ccdc78 KD (D) and Ccdc33 KD (E) embryos. Scale bar represents 50μm.

**Supplementary figure 8.**
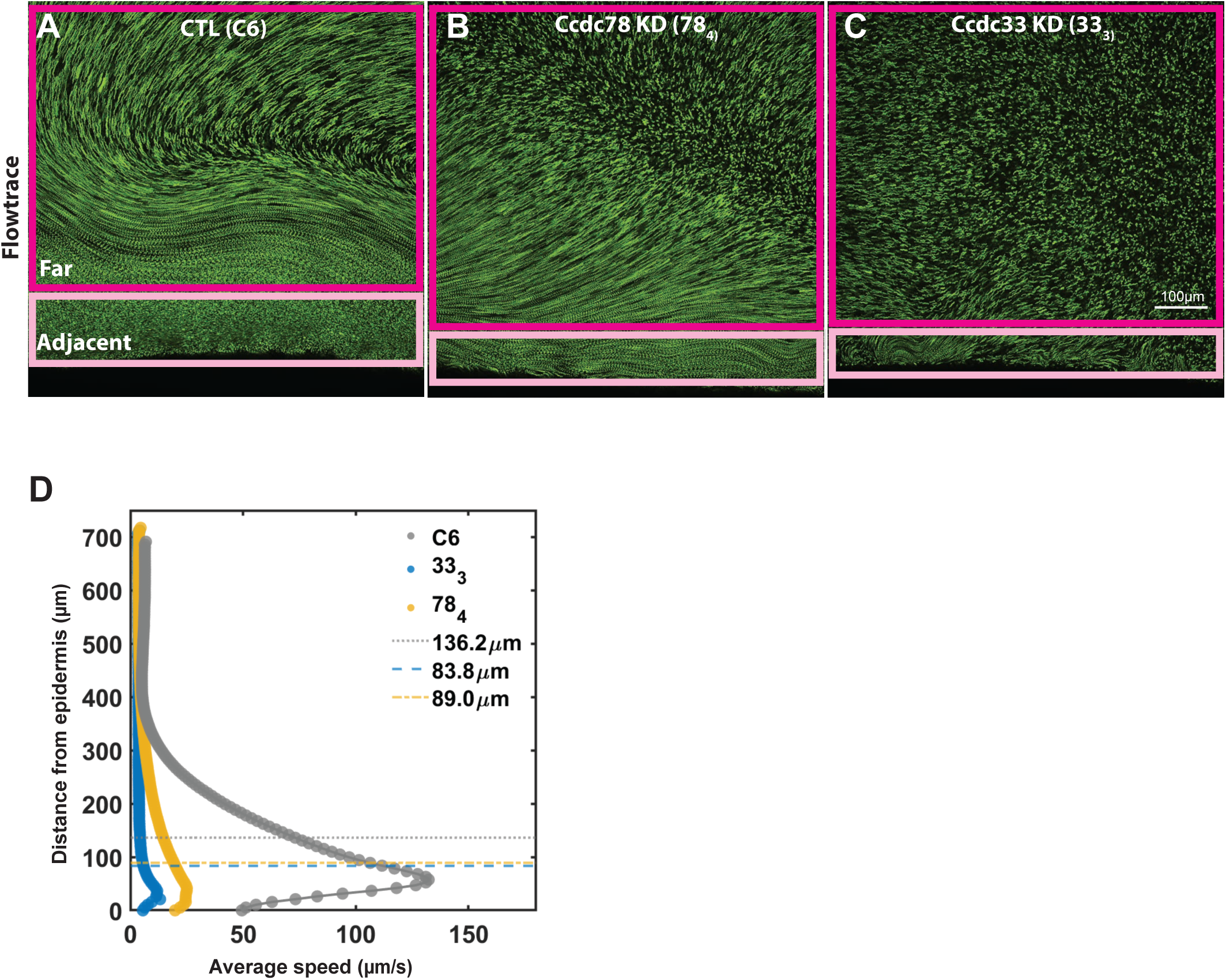
Bead flow analysis of control, Ccdc78 KD and Ccdc33 KD embryos. (A-C) 10seconds time averaged Flowtrace visualizations of one dataset each of control (A), Ccdc78 KD (B) and Ccdc3 KD (C). Far (magenta box) and adjacent (pink box) are highlighted with the box. The thresholds to define the adjacent region and the far region were determined to be the point of the steepest fall in flow speeds. (D) Peak speed averaged in control (black), Ccdc78 KD (magenta) and Ccdc33 KD (green). The bar graphs represent mean values, and the error bars represent standard error. (E) Average flow speeds plotted as a function of frame number. The time difference between each frame is 25mseconds. The purple dashed line corresponds to 22.5seconds and represents the total time duration of data used for analysis.

**Supplementary figure 9.**
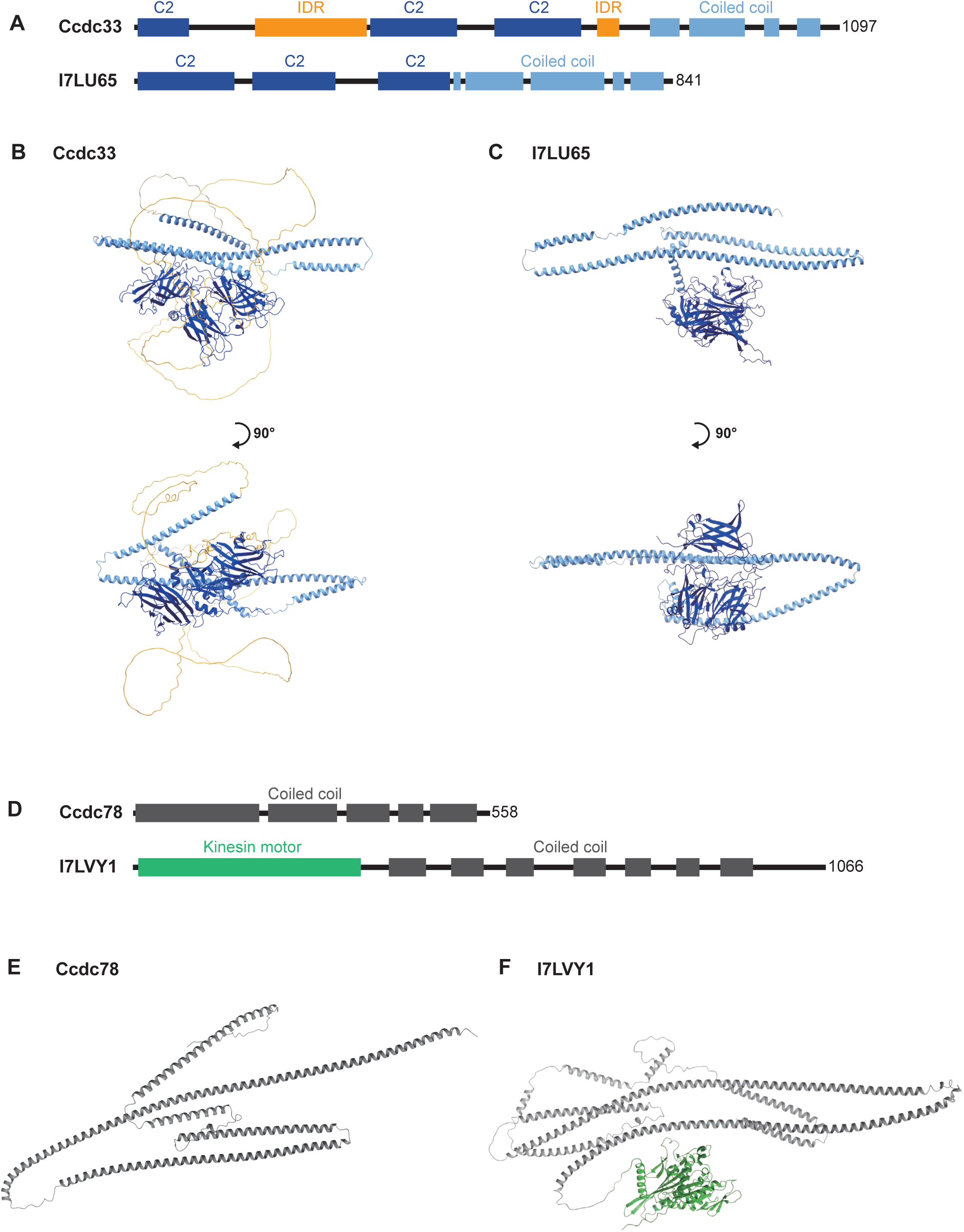
Structural differences of Ccdc78 and Ccdc33 in *Tetrahymena* and *Xenopus laevis*. (A) Structural domain of *Xenopus* Ccdc33 and *Tetrahymena* I7LU65. (B-C) Alphafold prediction of protein structures for *Xenopus* Ccdc33(B) and *Tetrahymena* I7LU65(C) with a 90-degree rotation. Coiled-coil regions are colored light blue, intrinsically disordered regions (IDRs) are colored yellow and C2 domains are colored dark blue. (D) Structural domain of *Xenopus* Ccdc78 and *Tetrahymena* I7LVY1. (E-F) Alphafold prediction of protein structures for *Xenopus* Ccdc78(E) and *Tetrahymena* I7LVY1(F). Coiled-coil regions are colored gray, and the kinesin motor domain is colored green.

**Supplemental Table 1:** APMS data for Ccdc78 Pulldowns. Raw data available on MassIVE https://massive.ucsd.edu/ProteoSAFe/static/massive.jsp (MSV000096822).

**Supplemental Table 2:** APMS data for Ccdc33 Pulldowns. Raw data available on MassIVE https://massive.ucsd.edu/ProteoSAFe/static/massive.jsp (MSV000096822).

**Supplemental Movie 1:** Multiciliated cell beating in control embryo in figure 9A.

**Supplemental Movie 2:** Multiciliated cell beating in Ccdc78 KD embryo in figure 9B.

**Supplemental Movie 3:** Multiciliated cell beating in Ccdc33 KD embryo with mild effect on metachrony and relatively uniform length of cilia, shown in figure 9C.

**Supplemental Movie 4:** Multiciliated cell beating in Ccdc33 KD embryo with stiffer cilia and severe defect in the metachronal wave, shown in figure 9D.

**Supplemental Movie 5:** Flowtrace movie of control embryo with fluorescent beads shown in Figure 9E.

**Supplemental Movie 6:** Flowtrace movie of Ccdc78 KD embryo with fluorescent beads shown in Figure 9F.

**Supplemental Movie 7:** Flowtrace movie of Ccdc33 KD embryo with fluorescent beads shown in Figure 9G.

## Acknowledgements

JBW is supported by the NHLBI (R01HL117164); JH was supported by the Korea Health Technology R&D Project through Korea. Health Industry Development Institute (KHIDI), funded by the Ministry of Health & Welfare, Republic of Korea (grant number: HI19C1095). EMM was supported by the NIGMS (R35GM122480), Army Research Office (W911NF-12-1-0390), and the Welch Foundation (F-1515). AH was supported by the (NHLBI R01HL173490). SB was supported by the NHLBI (HL146601, HL128370). TJP is supported by the Institute for Basic Science (IBS-R022-D1) and the Korean National Research Foundation (grant number: RS-2023-00225555). VNP was supported by startup funding from the University of Miami.

## Materials and Methods

### Animal husbandry and embryo manipulation

To induce the ovulation, female adult *Xenopus laevis* was injected with 400unit of hCG (Human chorionic gonadotropin) and incubated in 16℃ incubator for overnight. In vitro fertilization was performed by mixing the *Xenopus egg* with homogenized testis in 1/3X MMR (Marc’s Modified Ringer’s). Fertilized embryos were then de-jellied with 3% L-cysteine in 1/3X MMR (pH.7.8), washed and manipulated in 1/3X MMR.

### Plasmids, mRNA and Morpholino

Gene sequence for *Xenopus laevis* was obtained from Xenbase (https://www.xenbase.org/). Total RNA was purified from *Xenopus laevis* embryo and then reverse transcribed into cDNA library using M-MLV Reverse Transcriptase (Invitrogen). Coding sequence (CDS) of genes were amplified from the *Xenopus* cDNA library by PCR (Polymerase chain reaction) using Q5® High-Fidelity DNA Polymerase (NEB). Primer pairs were designed with restriction enzymes sites for cloning. Amplified genes and pCS10R vector containing fluorescence tag were digested with restriction enzymes, ligated together and inserted into competent cells by transformation. Cloned constructs were linearized to synthesize mRNA using mMESSAGE mMACHINE™ SP6 Transcription Kit (Invitrogen). The list of cloned genes are as follows: Armc9, Cep104, Ccdc78, Ccdc33, Spef1, Spag6, Cfap52, Ak7, Tsga10, EMTB.

Anti-sense morpholinos were designed to block RNA splicing based on the sequence from Xenbase database. The morpholinos were manufactured by Gene Tools. Ccdc78 and Ccdc33 morpholino sequences are as follows:

Ccdc78.L MO: 5’-CCCATTCCTTTCACTTACATTTTC-3’

Ccdc33.L MO: 5’-GGTCAGGTAGTCACAGTATAAGAA-3’

### Embryo microinjection and sample preparation

Embryo microinjection was done in 2% ficoll in 1/3X MMR. For the multi-cilia visualization, 2-cells of ventral-animal side in 4-cell stage embryos were injected with fluorescence tagged mRNA and morpholinos. For the nodal cilia visualization, two dorsal-ventral -cells in 4-cell stage embryos were injected with GFP-tagged Arl13b, BFP-tagged cent4 and mScarlet3-tagged Ccdc78 or Ccdc33.

For multi-ciliated cells live-imaging, *Xenopus* embryos were incubated until stage 26 after microinjection. Whole embryos were mounted between the cover glass with a small amount of 1/3X MMR and imaged immediately. For nodal cilia live-imaging, *Xenopus* embryos were incubated until stage 18 after the microinjection. Embryos were dissected laterally to dorsal posterior to obtain the gastrocoel roof plate (GRP) region. Dissected explants were mounted in the same way as whole embryos and imaged immediately.

### Image acquisition and analysis

Confocal images were acquired with Zeiss LSM700 laser scanning confocal microscope using a plan-apochromat 63X/1.4 NA oil objective lens (Zeiss). Structured illumination microscopy (SIM) images were acquired with Nikon DeepSIM microscopy using a 100X oil objective lens (Nikon).

Quantitative measurement of images was done using Fiji. The fluorescent intensity values of each measured pixel for each protein was normalized with the average intensity of the entire distal-most four microns of individual cilia. Graph generation and statistical analysis including error bars and mean ± SD and P values were performed using Prism 10 software.

### Cilia isolation and Negative staining for TEM

*Xenopus* embryos in tailbud stage were anesthetized with 0.05% Benzocaine in 1/3X MMR and washed with 1/3X MMR with several times to clear the debris from vitelline envelope and excess mucus. Embryos were transferred to the 2ml PCR tube and incubated with deciliation buffer (20mM HEPES pH 7.0, 112mM NaCl, 3.4mM KCl, 10mM CaCl2, 2.4mM NaHCO3, 20% EtOH, 1x protease cocktail) for 45seconds on nutator. The supernatant with isolated cilia was transferred to another 2ml PCR tube and centrifuged for 5 minutes in 13,000xg. The pellet was resuspended and fixed with 0.15% Glutaraldehyde for 10 minutes in room temperature. After centrifugation for 5 minutes in 13,000xg, the cilia pellet was resuspended and washed with 1X PBS, then centrifuged again for 5 minutes in 13,000xg. The final cilia pellet was resuspended with 20μl of 1X PBS.

For negative staining, 5μl of isolated cilia was placed on the grid (Formvar/Carbon 300 Mesh, Cu, Electron microscopy sciences) and sat for 1 minute. The grid was stained with 2% uranyl acetate solution and washed with water. Air-dried grids were stored in grid box for further imaging. The imaging of samples were done with JEOL NEOARM Low kV STEM.

### Protein purification and Immunoprecipitation

GFP-tagged Ccdc78 and Ccdc33 plasmid respectively were injected into 2-cell of ventral-animal side of the *Xenopus* embryos in 4-cell stage. Injected embryos were incubated until they reached stage 9, and their animal caps were dissected in Steinberg’s solution with gentamicin. When animal caps reached stage 26, animal caps were collected for protein extraction.

Protein purification and immunoprecipitation were performed with GFP-Trap agarose kit (Chromotek). Collected embryos were lysed with lysis buffer containing 1X protease inhibitor (Thermo Scientific™ Halt™ Protease Inhibitor Cocktail (100X), Thermofisher) and 1mM PMSF (Phenylmethylsulfonyl fluoride). Lysed samples were centrifuged in 4℃ for 15 minutes with 14000rpm to separate the lipid layer and collect clear lysates. GFP-agarose beads were equilibrated in dilution buffer according to the manufacturer’s protocol. GFP-agarose beads in dilution buffer were added to lysate and incubated at 4℃ for 1 hour. After protein binding, beads were centrifuged at 4℃ for 5 minutes with 2500xg. Pelleted beads were washed with wash buffer 2 times and added in 1.5X Laemmli sample buffer with 5% (v/v) 2-Mercaptoethanol. After boiling at 95℃, beads were pelleted with 4℃ centrifuge for 5 minutes and the supernatant was collected and stored at −80℃ for further analysis.

### Affinity purification Mass spectrometry

Immunoprecipitated proteins were resuspended in SDS-PAGE sample buffer and heated 5 min at 95°C before loading onto a 7.5% acrylamide mini-Protean TGX gel (BioRad). After 7 min of electrophoresis at 100 V the gel was stained with Imperial Protein stain (Thermo) according to manufacturer’s instructions. The protein band was excised, diced to 1 mm cubes and processed for in-gel trypsin digestion as in Goodman et al., 2018^74^. Digested peptides were desalted with 6µg-capacity ZipTips (Thermo Scientific), dried, and resuspended in 20µl of 5% acetonitrile, 0.1% acetic acid for mass-spectrometry. Peptides were separated using reverse phase chromatography on a Dionex Ultimate 3000 RSLCnano UHPLC system (Thermo Scientific) with a C18trap to EASY-Spray PepMap RSLC C18 column (Thermo Scientific, ES902) configuration eluted with a 3% to 45% gradient over 60 min. Spectra were collected on a Thermo Orbitrap Fusion Lumos Tribrid mass spectrometer using a data-dependent top speed HCD acquisition method with full precursor ion scans (MS1) collected at 120,000 m/z resolution. Monoisotopic precursor selection and charge-state screening were enabled using Advanced Peak Determination (APD), with ions of charge + two selected for high energy-induced dissociation (HCD) with stepped collision energy of 30% +/- 3% (Lumos) Dynamic exclusion was active for ions selected once with an exclusion period of 20 s (Lumos). All MS2 scans were centroid and collected in rapid mode. Raw MS/MS spectra were processed using Proteome Discoverer (v2.5) and the Percolator node to assign unique peptide spectral matches (PSMs) and protein assignments (FDR .01) to a X. laevis proteome derived from the 2023 UniProt Xenopus laevis reference proteome of 35,860 protein sequences with homeologs and highly related entries collapsed into EggNOG vertebrate-level orthology groups^75^. This database and the mass spectrometry data are available on MassIVE https://massive.ucsd.edu/ProteoSAFe/static/massive.jsp (MSV000096822).

In order to identify proteins significantly associated with each bait, we used the degust statistical framework (https://degust.erc.monash.edu/) to calculate both a log2 fold-change and an FDR for each protein enrichment based on the observed PSMs in the bait versus control pulldown. Settings used were “RUV (edgeR-quasi-likelihood), Normalization TMM, and Flavour RUVr” and at least 2 counts in at least 2 samples.

### RT-PCR

To confirm the efficacy of Ccdc78 and Ccdc33 morpholinos, morpholinos were injected into all cells at 4-cell stage of embryo. Total RNA was isolated with Trizol reagent at stage 26, and cDNA was synthesized with M-MLV reverse transcription kit (Invitrogen). Ccdc78, Ccdc33 and Odc1 were amplified by Taq-polymerase (Invitrogen) with the primers as follows:

Ccdc78 Forward: 5’-AGAAGATCGTGAGACCCCCC-3’

Ccdc78 Backward: 5’-TCAAGCTCTGCTGTGGCTTC-3’

Ccdc33 Forward: 5’-GCTTCCAGAACAACACCTACCTTT-3’

Ccdc33 Backward: 5’-GTGAGACAGGCGGAGAATCATCTA-3’

### Immunoblotting

*Xenopus laevis* embryos were injected with GFP-Ccdc78 or GFP-Ccdc33 in all cells at 4-cell stage with Ccdc33 or Ccdc78 morpholino, respectively. At stage 26, embryos were lysed in lysis buffer containing 1X protease inhibitor (Thermo Scientific™ Halt™ Protease Inhibitor Cocktail (100X), Thermofisher) and 1mM PMSF (Phenylmethylsulfonyl fluoride). Fat was removed by centrifuging the samples with 12000xg in 4℃ for 15 minutes. The clear lysate was added with 4X Laemmli sample buffer with 5% (v/v) 2-Mercaptoethanol and heated to 95℃ for 10 minutes. The protein samples were further processed for the SDS-PAGE and immunoblotting as described below.

Protein samples were loaded on SDS-PAGE and transferred to nitrocellulose membranes. The membrane was incubated in blocking solution (0.05% Tween-20 in TBS with non-fat powdered milk) at room temperature for 30 minutes. For immunoblotting, membranes were sealed with primary antibodies in 1% BSA solution for 1 hour at room temperature.

Secondary labeling was performed using horseradish peroxidase (HRP)-conjugated secondary antibodies for 1 h at room temperature. Chemiluminescence was performed with enhanced chemiluminescence substrate and imaged with Image Quant (LAS 4000, GE Healthcare).

The primary antibodies used were as follows:

mouse anti-beta-actin (1:10000, 66009-1, Proteintech); mouse anti-GFP (1:1000, sc-9996, Santacruz Biotechnology).

The secondary antibodies used were as follows:

Anti-mouse IgG HRP-conjugated (1:2000, 31430, ThermoFisher Scientific)

### Human ALI culture and immunostaining

Human trachea epithelial cells (HTECs) culture were prepared from airway epithelial cells isolated from tracheobronchial tissues as previously described^76^. Use of human tissues was reviewed by the Washington University Institutional Review Board. Anonymized human tissues were obtained from surgical excesses of deceased individuals and experimented by Code of Federal Regulations, 45 CFR Part 46, as not meeting criteria for human subject research. Briefly, epithelial cells were isolated from the tissues following incubation in pronase and differential adhesion on tissue culture plates. Basal cells were expanded in custom media, then released by trypsin treatment and cultured on Trans-well membranes (Trans-well, Corning, 3460). Cells were differentiated using air-liquid interface (ALI) conditions and media previously described^76^.

Cultured airway cells were immunostained in situ on the Trans-well membranes. Cells were fixed with 4% paraformaldehyde (PFA) for 10 min at room temperature. Non-specific antibody binding was blocked in buffer containing 3% bovine serum albumin and 0.1% TritonX100 in PBS for 1 hour at room temperature. Cells were then washed with 0.1% Tween20 in phosphate buffer saline (PBS) and incubated in primary antibodies diluted in blocking buffer. Primary antibodies were anti-CCDC78 (1:100, Proteintech, 26876-1-AP), anti-CCDC33 (1:100, Santacruz, sc-390852) and acetylated alpha tubulin (mouse, Clone 6-11B-1, 1:20.000; Sigma-Aldrich, Cat# T7451; RRID: AB_609894). After incubation at 4 °C overnight, the cells were then washed and incubated with species-specific, fluorescent-labeled secondary antibodies diluted in PBS for 45-60 min at room temperature. Membranes were mounted on glass slides in medium containing 4’, 6-diamidino-2-phenylindole (DAPI) to stain DNA (Fluoroshield, Sigma Aldrich, # F6057). Cells were imaged by wide-field fluorescent microscopy using an upright Leica 6000 microscope equipped with a cooled digital camera interfaced with imaging software (LAS X, Leica).

### Mouse tissue immunostaining

The trachea and oviduct were collected from P21 mouse. Formalin-fixed and paraffin-embedded (FFPE) tissues were sectioned at 4 µm. The FFPE sections were dewaxed, rehydrated, and boiled in antigen retrieval solution Histo-VT-One (Nacarai). After blocking with Blocking-One-Histo (Nacarai), the sections were incubated with anti-CCDC78 antibody (1:100, Proteintech, 26876-1-AP) and acetylated-tubulin (1:300, Proteintech, 66200-1-Ig) overnight at 4 °C, then with Alexa Fluor-conjugated secondary antibodies (1:500, Invitrogen, A-21206, A-21203). The sections were mounted in Fluorescence-Mounting-Medium (Dako). Immunofluorescence imaging was performed on a LSM800 confocal laser scanning microscope (Carl Zeiss).

### *In vitro* microtubule bundling

*In vitro* translation of GFP, GFP-Ccdc78 and GFP-Ccdc33 proteins were performed using TnT® Coupled Wheat Germ Extract System (Promega). Briefly, 2μg of CS10R-GFP, CS10R-GFP-Ccdc78 and CS10R-GFP-Ccdc33 plasmids were incubated with 30μl of TnT® SP6 High-Yield Wheat Germ Master Mix in total 50μl reaction, 25℃ 2 hours. The concentration of proteins was determined with western blot analysis by comparing translated proteins to serial dilutions of GFP peptide.

To polymerize microtubules *in vitro*, 2μl of 647-labeled tubulins (2mg/ml)(Pursolutions, 064705) were diluted in 6μl of 1mM GTP containing tubulin buffer (80mM PIPES pH 6.9, 1mM EDTA, 1mM MgCl2, 10% glycerol). Tubulins were polymerized for 1 hour in 37℃ water bath and stabilized by adding 35μM of Taxol solution to the polymerized microtubules. After an additional 20 minutes incubation in 37℃ water bath, microtubules were stored at room temperature covered with foil to avoid light.

For microtubule bundling assay, stabilized microtubules were diluted into tubulin buffer in 1:20 dilution. 6μl of diluted microtubules were mixed with 4μl GFP, 4μl GFP-Ccdc78, 4μl GFP-Ccdc33 and 2μl GFP-Ccdc78 + 2μl GFP-Ccdc33. After incubation at room temperature for 20 minutes, 10μl of each mixture was gently squashed between slide glass and coverslip and sealed with VALAP (1:1:1 Vaseline: lanolin: paraffin wax).

### Bead flow analysis

*Xenopus* embryos were grown up to stage 25-26 and mounted onto an imaging chamber in 1/3X MMR with green fluorescent beads (Fluospheres^TM^ polystyrene, 1.0μm, yellow-green 505/515). The bead flow movies and live cilia beating videos were taken on a Nikon W1 Spinning Disc Confocal at an image rate of 40 frame per seconds and 80 frame per seconds respectively.

Fluid flow quantification was carried out by processing a total of 18 videos with tracer particles live-imaged at 40fps for 30s duration in each case. The datasets included 6 videos each for control, 78KD and 33 KD. The fluid flow fields were visualized using the Flowtrace plugin in ImageJ ^53^ with a time window of 10s. Further, the timelapse image sequences were analyzed by implementing the Particle Image Velocimetry (PIV) technique using the MATLAB-based

PIVlab package^54^. The velocity vectors were computed between consecutive timepoints for all the datasets using individual frames extracted from the videos (40fps, 30s total duration). For all the frames, the epidermis at the bottom of images was masked, and the images were pre-processed using the contrast limited adaptive histogram equalization (CLAHE) option (window size 16 pixels). The velocity vectors were computed via the FFT cross-correlation method with a first pass on 64x64 pixels interrogation window and the second pass on 32x32 pixels interrogation window with 50% overlap. Velocity vector validation was performed using appropriate velocity limits, standard deviation filters and interpolation of missing vectors.

The velocities in the directions, parallel to the epidermis and perpendicular to the epidermis, and the resultant flow speeds (velocity magnitude) were extracted at each time point. The results were calculated from the first 22.5s of the videos (first 900 frames) for all datasets, and the remaining duration with fluctuations due to larval twitching was discarded. The fluid flow were quantified by comparison of the flow speeds averaged over time, across the area adjacent to the epidermis and far from the epidermis. The thresholds to define the adjacent region and the far region were determined to be the point of the steepest fall in flow speeds (location of minimum velocity gradient) from the epidermis. These thresholds were computed from the average speed profiles in the direction perpendicular to the epidermis, calculated by averaging flow speed over time and in the direction parallel to the epidermis. The peak and minimum flow speeds were also calculated from the average speed profile.

### Alphafold3 structure prediction

The protein structure predictions were predicted by Alphafold3 using Alphafold server (https://alphafoldserver.com/).

